# Major trends and environmental correlates of spatiotemporal shifts in the distribution of genes compared to a biogeochemical model simulation in the Chesapeake Bay

**DOI:** 10.1101/2023.01.09.523340

**Authors:** Sarah Preheim, Shaina Morris, Yue Zhang, Chris Holder, Keith Arora-Williams, Paul Gensbigler, Amanda Hinton, Rui Jin, Marie-Aude Pradal, Anand Gnanadesikan

## Abstract

Microorganisms mediate critical biogeochemical transformations that affect the productivity and health of aquatic ecosystems. Metagenomic sequencing can be used to identify how the taxonomic and functional potential of microbial communities change in response to environmental variables by investigating changes in microbial genes. However, few studies directly compare gene changes to biogeochemical model predictions of corresponding processes, especially in dynamic estuarine ecosystems. We aim to understand the major drivers of spatiotemporal shifts in microbial genes and genomes within the water column of the Chesapeake and highlight the largest discrepancies of these observations with model predictions. We used a previously published shotgun metagenomic dataset from multiple months, sites, and depths within Chesapeake Bay in 2017 and a metatranscriptomic dataset from 2010-2011. We compared metagenomic observations with rates predicted with a comprehensive physical-biogeochemical model of the Bay. We found the largest changes in the relative abundance of genes involved in carbon, nitrogen, and sulfur metabolism associated with variables that change with depth and season. Several genes associated with the largest changes in gene abundance are significantly correlated to corresponding modeled processes. Yet, several discrepancies in key genes were identified, such as differences between genes mediating nitrification, higher than expected abundance and expression of denitrification genes in aerobic waters, and nitrogen fixation genes in environments with relatively high ammonia but low oxygen concentrations. This study identifies processes that align with model expectations and others that require additional investigation to determine the biogeochemical consequences of these discrepancies and their impact within an important estuarine ecosystem.

## Introduction

Microorganisms are responsible for many biogeochemical processes that influence the health and productivity of estuarine and coastal aquatic ecosystems, so studying the factors that influence their abundance and activity is important. Many microbially mediated processes are beneficial to the ecosystem, such as photosynthesis and organic matter degradation and recycling (Canfield et al. 2005), while other processes are noxious or detrimental, such as harmful algal blooms (Griffith and Gobler 2020), oxygen consumption resulting in hypoxia (Kemp et al. 2009), and greenhouse gas production (Holmquist et al. 2018). Because of their importance to the ecosystem, it is critical to understand relationships between microbial processes and environmental variables, so they can be accurately represented in models for management and planning purposes.

Metagenomic analysis could provide insight into the microbial community mediating key biogeochemical processes influencing oxygen demand, nutrient cycles, or greenhouse gas production (Riesenfeld et al. 2004). Metagenomic analysis has the potential to be more sensitive at identifying certain biogeochemical processes than chemical signatures alone (Reed et al. 2014). Researchers have used genetic observations to constrain the structure of biogeochemical models relatively successfully (Coles et al. 2017; Louca et al. 2016a; Preheim et al. 2016; Reed et al. 2014), although typically this has been done in ecosystems assumed to be at or near steady-state. In some cases, there seems to be a strong relationship between the observed genes and the function they mediate, such as with the denitrification gene *nirS* and nitrous oxide production in the Chesapeake Bay (Ji et al. 2018). In other cases, this relationship breaks down, even for the same *nirS* genes (Bowen et al. 2014). More work is needed to understand factors that influence changes in microbial community gene composition in space and time and how these changes may be used to inform biogeochemical models.

Gene abundances may provide a measure of the potential for the community to make enzymes that mediate the associated biogeochemical transformations. In some cases, this potential corresponds well with where and when these transformations occur, while other times this correspondence can break down. Because many of the genes selected for analysis are metabolic genes associated with obtaining energy for growth and cellular maintenance, the expectation is that cells carrying these genes will reproduce in environments with sufficient energy available and nutrients to support these processes. However, a meta-analysis of the relationship between gene abundance or expression and measured process rates demonstrated a significantly positive correlation only 38% of the time (Rocca et al. 2015). There are many reasons why the observed relationship between genes and processes could break down, such as by recent migration, disequilibrium between the microbial community and the environment, phenotypic plasticity, functional redundancy, or dormancy (Hall et al. 2018). Hall and colleagues (Hall et al. 2018) argue that a robust conceptual framework is needed to link the resident microbes to the processes they mediate. Although a few studies have been conducted within aquatic ecosystems to link microbial populations, genes, transcripts and proteins to the observed changes in chemical concentrations in these systems (Arora-Williams et al. 2022; Haas et al. 2021; Louca et al. 2016a; Preheim et al. 2016; Reed et al. 2014), much work remains to be done to understand how such linkages vary across ecosystems. As we continue to map genes to their ecosystems using a framework that provides a mechanistic understanding of the factors that influence these processes, our understanding of these relationships will continue to improve and lead to further discoveries

The Chesapeake Bay (hereafter, “the Bay”) is an important example of a critical coastal ecosystem that has experienced significant declines in ecosystem health and productivity associated with nutrient and sediment pollution. It supports over a billion dollars in iconic fisheries such as blue crabs, oysters and rockfish (National Marine Fisheries Service 2018), but was even more productive in pre-colonial days before the watershed was significantly impacted by human activity. Changes in the watershed were associated with shifts in productivity from benthic diatoms and seagrasses to near-surface phytoplankton and decreases in the abundance of oysters (Rothschild et al. 1994), blue crabs (Rugolo et al. 1998) and various species of commercially valuable fish. One driver of these species declines has been the increase in hypoxia each summer in significant portions of the Bay (Da et al. 2018), with occasional episodes of anoxia or euxinia (Cornwell and Sampou 1995). Sediment loads and nutrient pollution are two main drivers of oxygen depletion. Both have increased in response to changes associated with urbanization and draining of wetlands (Pasternack et al. 2001), poor wastewater and stormwater management [e.g. (Fisher et al. 2021)], and increases in agricultural lands within the watershed (Brush 2009).

Because water quality in the Chesapeake Bay has been impacted by human activity, efforts have been made to reduce sediment and nutrient pollution to improve water quality. The District of Columbia and states of Maryland, Virginia, and Pennsylvania established the Chesapeake Bay Program under Federal auspices to lead and direct efforts to restore the Bay (https://www.chesapeakebay.net/who/bay_program_history, last access 8/24/2022). The Chesapeake Bay Program represents a major effort by states and the Federal government to restore the Bay and includes a substantial monitoring program. While this program has aided in the development of several models (Cerco and Noel 2017; Da et al. 2018; Testa et al. 2018; Testa et al. 2014) that have significant skill in predicting interannual variability in hypoxia (Irby et al., 2017), much of this skill appears to derive from their ability to capture changes in mixing and ventilation (Scully 2016). There are indications that this approach may be insufficient for predicting changes in hypoxia associated with nutrient reduction, as there is evidence each unit of nutrient input is producing more hypoxia now than in times past (Hagy et al. 2004; Kemp et al. 2005; Murphy et al. 2011; Scavia et al. 2021). This makes it difficult to predict how effective further nutrient reductions will be in driving improved oxygen concentrations over time. As a result, the latest plan for the Bay program looks to “identify key missing processes and update the code to address knowledge gaps as they are filled” https://www.chesapeakebay.net/what/programs/modeling/phase_7_model_development, last access date 8/24/22). Our goal in this manuscript is to examine whether metagenomics can help with this important objective.

The response of the microbial community to changes in nutrients, sediments and the environment could affect the recovery trajectory of the Bay, potentially resulting in hysteresis with a delayed response to improvements in water quality (Middelburg and Levin 2009). Previous work in the Bay has identified major drivers of community changes and abundant and active taxa and functions that were under-appreciated within this ecosystem. Seasonal changes have been identified as the most significant driver of changes in community composition (Arora-Williams et al. 2022; Kan et al. 2006; Wang et al. 2020), with shifts associated with spatial distribution using 16S rRNA gene sequencing (Arora-Williams et al. 2022; Wang et al. 2020) along the length of the Bay being of secondary importance. Community composition (Crump et al. 2007) and gene expression (Hewson et al. 2014) change in response to low oxygen conditions, with specific taxa and functions predictive of hypoxic conditions (Arora-Williams et al. 2022). Previous work identified shifts in community composition under sulfidic conditions (Crump et al. 2007), with expression of genes associated with sulfate reduction increasing in deep anoxic water (Hewson et al. 2014). Through analysis of taxa and genome composition from metagenome assembled genome analysis, sulfide oxidation coupled to at least partial denitrification was identified as a common microbial response to low oxygen in the water column that was previously under-appreciated within this ecosystem (Arora-Williams et al. 2022). Metagenomic analysis could identify other currently under-appreciated microbially mediated processes or reveal delayed or accelerated on-set of specific processes that could have a substantial influence on nutrient cycles, oxygen concentrations, or greenhouse gas emissions.

In this paper, we examine the primary patterns of spatiotemporal variation in genes taken from 38 samples collected in Chesapeake Bay during the spring and summer of 2017, comparing observations with model predictions. We focus on a subset of genes that are involved in photosynthesis, aerobic metabolism, and anaerobic metabolism. We find that many genes shift as expected in relation to environmental parameters such as temperature and nutrients. However, several results emerge from this analysis that are not expected based on the model predictions. First, genes associated with denitrification and dissimilatory reduction of nitrate to ammonia (DNRA) are abundant and expressed in relatively well oxygenated waters during the spring. Nitrogen fixation genes are also abundant and expressed in locations with relatively high levels of ammonia, which was not included in the model. Anoxygenic photosynthesis genes were much more common than expected and found in oxygenated water. Finally, more work is needed to elucidate where, when, and who is involved in nitrification within this ecosystem. Future work should focus on determining whether these processes have a significant impact on Bay-wide nitrogen cycling, and if so, whether they should be included in models.

## Methods

### Description of metagenomic and metatranscriptomic datasets

We used shotgun metagenomic sequence data from our previously published dataset [BioProject PRJNA612045, (Arora-Williams et al. 2022)]. Our previous publication focused on the 16S rRNA gene analysis across a larger set of samples, for which we conducted a limited analysis of the shotgun metagenomic data to confirm population functional capabilities and activity. Here, we focus on the analysis of the shotgun metagenomic data itself, focusing on metabolic genes to compare to a biogeochemical model. Briefly, water samples for metagenomic analysis were collected in April, June, July, and August in 2017 mainly from stations CB3.3C (near the Bay Bridge), CB4.4, CB5.3, CB6.2, and CB7.3 (near the Atlantic Ocean) as part of the Eyes on the Bay water quality monitoring program. Samples were also collected from stations CB4.2C and CB4.3C in April and July. Samples for each month were collected over the course of up to one week. At each station, water from 1 meter from the surface and 1 meter from the bottom was pumped through tubing into a 50 ml conical tube. Samples were placed immediately in a freezer on the boat and stored at −20 °C for up to 3 months. Water was thawed in 32 °C water bath for 5 minutes, filtered through a 0.2 *μ*m filter, and stored at −80 °C for up to a year before processing. A suite of environmental variables measured for each sample was obtained from the Chesapeake Bay Foundation Data Hub (https://datahub.chesapeakebay.net/, Last access date 08/01/21). In this paper we consider the following subset of these variables: temperature, salinity, dissolved oxygen, nitrate, nitrite, ammonium, phosphate, dissolved organic nitrogen and phosphorus, particulate nitrogen and phosphorus, chlorophyll, phaeopigments and pH. A list of stations, locations, dates, and environmental conditions are listed Table S1.

In a separate effort from the Eyes on the Bay program, samples were collected at one site (CB3.3C) at 0, 4, 8, 10, 12, 14, and 20 m depth on June 15, July 15, and Aug. 14, 2017. A peristaltic pump was used to filter water through a 0.2 *μ*m filter, which was placed on dry ice on the boat and stored in a −80 °C freezer for up to a year before processing. We have previously demonstrated that differences in sample storage (i.e. freezing the filter on the boat or freezing water and filtering in the lab) does not significantly change the microbial community composition (Arora-Williams et al. 2022). This separate dataset was only used to verify the distribution of genes in the water column, but was not used for other parts of the analysis because comparable environmental measurements (e.g. ammonia, nitrate, etc.) were not available.

A metatranscriptomic dataset from a previous study (Hewson et al. 2014) was also used to determine whether there was evidence for expression of genes of interest at certain times of the year or positions in the water column. Sequences were retrieved from NCBI (BioProject PRJNA222777) and additional files missing from the NCBI dataset were also provided to us by the authors (Table S2).

### Shotgun metagenomic library prep, sequencing and analysis

DNA was extracted with DNeasy PowerWater Kit (Qiagen) following the manufacturer’s protocol. Shotgun metagenomic libraries were prepared with the Illumina Nextera Flex kit following the manufacturer’s protocol and sequenced on an Illumina HiSeq at the JHU’s Genetic Research Core Facility. For this analysis, poor quality bases were trimmed with Trimmomatic (Bolger et al. 2014) and assembled with Spades (Bankevich et al. 2012) using the --meta option for metagenomic analysis. Genes were identified with Prodigal (Hyatt et al. 2010) and classified with GhostKoala (Kanehisa et al. 2016) using the entire database (genus_prokaryote + family_eukaryote + viruses).

Custom scripts were used to determine relative gene or transcript abundance by mapping reads to the assembly and calculating a scaled relative abundance metric similar to transcripts per kilobase million (TPM), where each library is normalized to 10^6^ reads and transcripts mapping to each gene are normalized to a 1 kb gene length. Bowtie2 (Langmead and Salzberg 2012) was used to map reads to the assembled contigs, using only those with high quality (-q 20), and samtools (Danecek et al. 2021), bedtools (Quinlan and Hall 2010), and custom perl scripts were used to determine the TPM for each gene. TPM was calculated by first determining reads per kilobase (RPK), as 1000 times the number of reads mapped to the gene divided by gene length. Next, reads per kilobase million (RPKM) was calculated as 10^6^ times RPK divided by the total number of mapped reads. Finally, transcripts per kilobase million (TPM) is calculated for each gene (*i*) in a library of *n* genes as 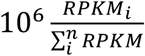. Scripts and code used to analyze this data can be found at https://github.com/spacocha/ChesapeakeBay_shotgun_metagenomics.

Four negative control samples and three samples re-sequenced from CB3.3C station separately from the water quality monitoring program were included in the library preparation, sequencing, assembly, and read mapping, but were excluded from the rest of the analysis. Of the 41 experimental samples, two samples resulting in less than 1,000 reads and one sample whose provenance could not be verified were excluded from further analysis, resulting in 38 total samples. After trimming and quality control, our dataset resulted in just over 15 million protein coding genes. Just under 4 million protein coding genes (26.1%) could be classified using the Kyoto Encyclopedia of Genes and Genomes (Kanehisa et al. 2017), with 17046 individual assignments. Approximately 3.5 million (89.2%) could be assigned a taxonomic classification.

### Description of key metabolic gene list

The KEGG database hierarchy was used to identify genes involved in key metabolic processes. We chose 76 genes involved in oxygenic and anoxygenic photosynthesis, assimilatory and dissimilatory nitrate reduction to ammonia, denitrification, nitrification, nitrogen fixation, assimilatory sulfate reduction, dissimilatory sulfate reduction and oxidation, SOX cycling, carbon fixation and methane cycling. We looked for but did not find the genes coding hydrazine synthase (*hzs*) and hydrazine dehydrogenase (*hdh*), which are involved in anaerobic ammonia oxidation. Finally, we chose a set of 22 housekeeping genes that were shared between the set of single-copy genes used for validation of bacteria and archaeal genomes in a previous study (Ishii et al. 2013). These genes were found to have low variance across samples (standard deviation of the log abundances were generally less than 0.25) and mean normalized TPM values were around 280 (± 80). Key metabolic genes and single-copy house-keeping control genes are listed in Table 1.

**Table 1:** List of chemical cycles (left column), pathways associated with these cycles (center column) and associated genes (right column) used in this analysis.

| Process | Pathway | Gene abbreviation<sup>1</sup> |
| --- | --- | --- |
| Oxygenic photosynthesis | Photosystem I | *psaA,B,C,D,E,F* |
|  | Photosystem II | *psbA,B,C,D,E,F,O* |
|  | Photosynthetic Electron transport | *petA,B,C,D,E,F,J,N* |
| Anoxygenic photosynthesis | Anoxygenic Photosystem II | *pufL,M* |
| Nitrogen cycling | Assimilatory nitrate reduction | *narB,NR,nasA,NIT6,nirA* |
|  | Dissimilatory nitrate reduction to ammonia | *narG,Z/nxrA, narH,Y/nxrB, narI,V* |
|  | Denitrification | *napA,B,nrfA,H narG,Z/nxrA, narH,Y/nxrB, narI,V, nirK,S, napA,B, norB,C, nirB,D,nosZ* |
|  | Nitrification | *pmoA/amoA, pmoB/amoB, pmoC/amoC, hao, narG,Z/nxrA, narH,Y/nxrB* |
|  | Nitrogen fixation | *nifD,K,H* |
|  | Anaerobic ammonia oxidation | ***Hzs<sup>2</sup>, hdh*** |
| Sulfur Cycling | Assimilatory sulfate reduction | *Sat,cysI,J, sir* |
|  | Dissimilatory sulfate reduction and oxidation | *Sat,aprA,B, dsrA,B* |
|  | Thiosulfate oxidation | *soxA,B,C,D,X,Y,Z* |
| Oxygenic photosynthetic carbon fixation | Rubisco | *rbcL,S* |
| Methane cycling | Methanogenesis | *mcrA,B,C,D,G, hdrA2, hdrB2* |
|  | Methane oxidation | *pmoABC, mmoBCD, mmoXYZ, mdh, mxaACDGKL, xoxF* |
| Housekeeping | Single-copy housekeeping genes | *IARS, AARS, MARS,DARS, PARS,RARS, FARSB, HARS, pyrG, ksgA,nusG, RP-L1,10, 11,13, 14,15,18, 2,22, 3* |

<sup>1</sup>KEGG ID for gene names of metabolic genes found in SI; <sup>2</sup>Genes in bold were sought in the dataset but not found.

Certain genes were characterized beyond their initial classification. *DsrAB* genes were classified as oxidative or reductive types by comparing the sequences to a dataset of oxidizing and reducing forms published previously (Muller et al. 2015), assuming the type from the best BLAST hit using blastn (Altschul et al. 1990). Viral *amoC* genes identified in a previous study (Ahlgren et al. 2019) were downloaded from JGI and compared to *amoC* genes in this dataset by BLAST. Taxonomic classification was obtained from GhostKoala and only used for genes that were given a KO number.

### Empirical orthogonal functions (EOF) analysis of gene abundance

TPM values were log transformed, with zero values set to 0.01. This produces a roughly normal distribution for the genes in Table 1, allows us to look at relative changes in genes, and ensures that the patterns we find are not dominated by changes in a few, relatively abundant genes. Anomalous abundances were computed by removing the mean log gene abundance for each gene. These were then used to compute empirical orthogonal functions (EOFs) yielding a set of spatiotemporal distributions of covarying genes that capture the largest modes of variation in relative abundance. Genes that appear in the same EOF with similar loadings will then change by similar factors across stations, irrespective of their mean abundance. Genes that appear in the same EOF with loadings that have similar magnitude but opposite signs will change in opposite directions (i.e. when the abundance of one doubles the abundance of the other will go down by a factor of 2). EOFs are computed by taking the singular vector decomposition of the matrix of anomalous abundances and represent eigenvectors of the covariance matrix. The loadings on each EOF were used to identify co-variance of key genes. We also generated analyses using powerlaw transformations of the gene abundance. As these analyses yielded almost identical patterns of variability for the top two EOFs we do not present them in detail.

When analyzing relationships between particular biogeochemical processes and hydrographic parameters we also combined relative gene abundances across multiple genes associated with these processes. We focus primarily on genes/processes where more than 50% of the observed variability is captured by a single EOF. We used stepwise linear regression to find the variables that explained the largest fraction of the variance. We performed regressions on both the spatiotemporal pattern associated with the EOFs and individual genes using either the top two or three variables depending on whether we were trying to compare predictors that were common to both models and observations, or simply trying to get the best fit to the data using observational predictors.

### Description of the biogeochemical model

In response to our population-level observations from a previous paper (Arora-Williams et al. 2022), we sought to create a working model of the Chesapeake Bay which includes additional nitrogen and sulfur cycling processes which are missing from many Bay models. This model is a coupled physical-biogeochemical model described in Jin et al. (Jin et al. 2022). The biogeochemical component of this model combines the ChesROMS_ECB model of Da et al. (Da et al. 2018), which was developed to simulate coupled nitrogen, oxygen and carbon cycling within Chesapeake Bay, with the RedoxCNPS model of Al-Azhar et al. (Al Azhar et al. 2014) which was developed to simulate cryptic sulfur cycling in the open-ocean oxygen-minimum zone off Peru. Both of these codes are descendants of that developed by Fennel et al. (Fennel et al. 2006). The model simulates the transport and interaction of ten dissolved inorganic species (nitrate, nitrite, ammonia, phosphate, sulfate, hydrogen sulfide, oxygen, dissolved inorganic carbon, total alkalinity), five dissolved organic species (semilabile and refractory dissolved organic nitrogen and carbon, semilabile phosphorus), six particulate species (small and large particulate N, P and C which sink at different rates) and one class each of phytoplankton and zooplankton. Apparent oxygen utilization is computed from dissolved oxygen concentration and the saturation concentration of oxygen (a function of temperature and salinity).

Phytoplankton productivity in this model is limited by nitrogen, phosphorus and light (though primarily by light throughout the year and nitrogen in the summer) and modulated by temperature. Production is primarily balanced by the exudation of semi-labile DOM and coagulation into large and small detrital particles, with faster and slower sinking velocities respectively. A small fraction of phytoplankton is grazed by zooplankton, but this term represents an insignificant part of the overall budget. The presence of DOM and particulate matter creates a demand that can be supplied by oxygen, nitrate (when oxygen is not available) or sulfate (when neither oxygen or nitrate is available). The respiration of organic matter using sulfate produces hydrogen sulfide, which can be oxidized by oxygen, nitrate, or nitrite. Additionally, anaerobic ammonia oxidation (annamox) can act as a loss term for fixed nitrogen. Some fraction of dissolved organic material hitting the bottom is resuspended, some is buried, and the remainder is assumed to remineralize to ammonia within the sediments with oxygen, nitrate or sulfate serving as potential oxidizers. In the absence of light and presence of oxygen, ammonia is oxidized first to nitrite, then to nitrate. The model thus implicitly or explicitly includes photosynthesis, carbon fixation, nitrification, heterotrophic denitrification, sulfate reduction, and sulfide oxidation.

This model is embedded within the ChesROMS physical model as described by Feng et al. (Feng et al. 2015). The model simulates the advection and mixing of water within the Bay as modulated by observed winds, tides, surface heat and moisture fluxes and riverine inputs. The model utilizes terrain-following coordinates with 20 levels in the vertical and has variable horizontal resolution, with a nominal resolution of about 1km at the stations we will be examining. The model shows a skill in simulating oxygen comparable to the model of Da et al. (Da et al. 2018) which has been used by the Virginia Institute of Marine Sciences to nowcast hypoxia within the Bay, as well as a significant number of other variables (Fig. S1). Model outputs are presented as daily averages.

### Gene abundance relationships with observed and modeled data

Log normalized TPMs were correlated with observed or modeled data using corr function in MATLAB. A set of random variables was generated and correlated with all abundant genes, along with observed environmental variables (oxygen, ammonia, phosphate) and modeled processes (sulfide oxidation) shown in the figures. Correlations with absolute values above 0.41 are significant at p < 0.01. Many genes show correlations with environmental variables that lie above this level-for example 17% of genes have correlations with temperature that lie outside the range [−0.41 0.41]. Looking across several environmental variables (Fig. S2), only about 5% of all genes exhibited correlations above 0.5, except for phosphate, where correlations above 0.54 were in the top 5%. Thus, we interpret correlations between genes with the associated observed or modeled variable above 0.5 as unlikely to arise by chance and to be within the top 5% of all such correlations.

## Results

### Key genes and taxa show substantial variation across the dataset

There were several major trends driving a substantial amount of variability in the abundance of key genes and taxa in this dataset. Changes in the measured environmental variables over the course of the spring and summer of 2017 were typical for the Bay, including substantial increases in temperature and phosphate and decreases in oxygen and pH below the pycnocline during the summer (Fig. S3). The pattern of taxonomic changes (Fig. 1a) occurring in space and time within this dataset was generally similar to taxonomic changes previously identified by 16S rRNA gene survey of the same samples (Arora-Williams et al. 2022), with a correlation of 0.67 between the metagenomic and 16S rRNA gene relative abundance datasets across the 14 most abundant phyla and Proteobacteria sub-phyla (Fig. S4). Some obvious and expected trends include more Cyanobacteria in surface samples than bottom water samples (Fig. S5), and Actinobacteria decreasing and Alphaproteobacteria increasing in relative abundance towards the mouth of the Bay in surface samples (Fig. S6).

**Figure 1.**
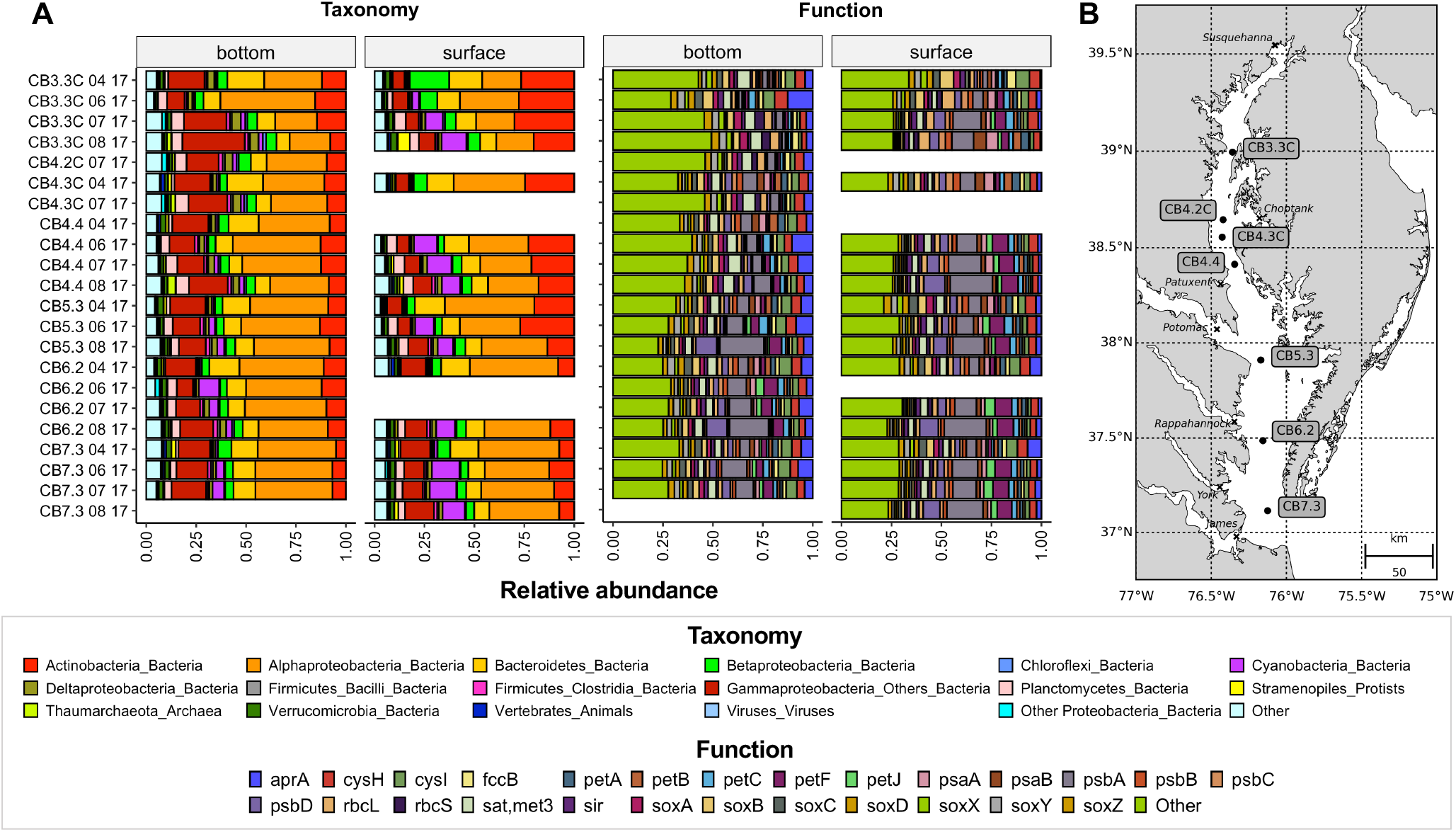
Location of sampling stations and changes in taxa and function within surface and bottom samples in space and time in the Chesapeake Bay. (A) Changes in relative abundance of taxa and functions predicted from the metagenomic data for various stations and dates. Bars represent the percent of all classified taxa or functions (unclassified taxa and genes are not shown). Samples are grouped into bottom or surface water samples and arrayed by station (CB3.3C-CB7.3) and date (MM YY). (B) Map of the Chesapeake Bay with sampling stations labeled.

Although there was limited variability across samples when considering all genes assigned functional categories in the KEGG database (Fig. S7), more substantial changes in space and time (Fig. 1a) were observed in a set of 76 key metabolic genes involved in photosynthesis, aerobic metabolism, and anaerobic metabolism (Table 1). Some obvious and expected trends include a higher relative abundance of photosynthetic genes (e.g. *psbA*) in the surface as compared with bottom water samples (Fig. S8) and more genes involved in sulfur oxidation and/or reduction (e.g. *aprA*) in the bottom (Fig. S9).

### Major modes of variation in metabolic genes are predictable from measured environmental parameters

To further determine factors associated with the largest spatiotemporal changes in the 76 metabolic genes outlined in Table 1, we used the top four empirical orthogonal functions (EOFs) to identify the major changes in gene relative abundance with depth and time (Fig. 2). The top two EOFs found with this subset of genes are clearly related to the top three EOFs when examining a larger subset of the most abundant genes (8,714 of the most abundant genes detected at more than 80% of the sites), demonstrating that these patterns are connected to major drivers of changes in gene composition across the entire dataset (see SI for more information).

**Figure 2.**
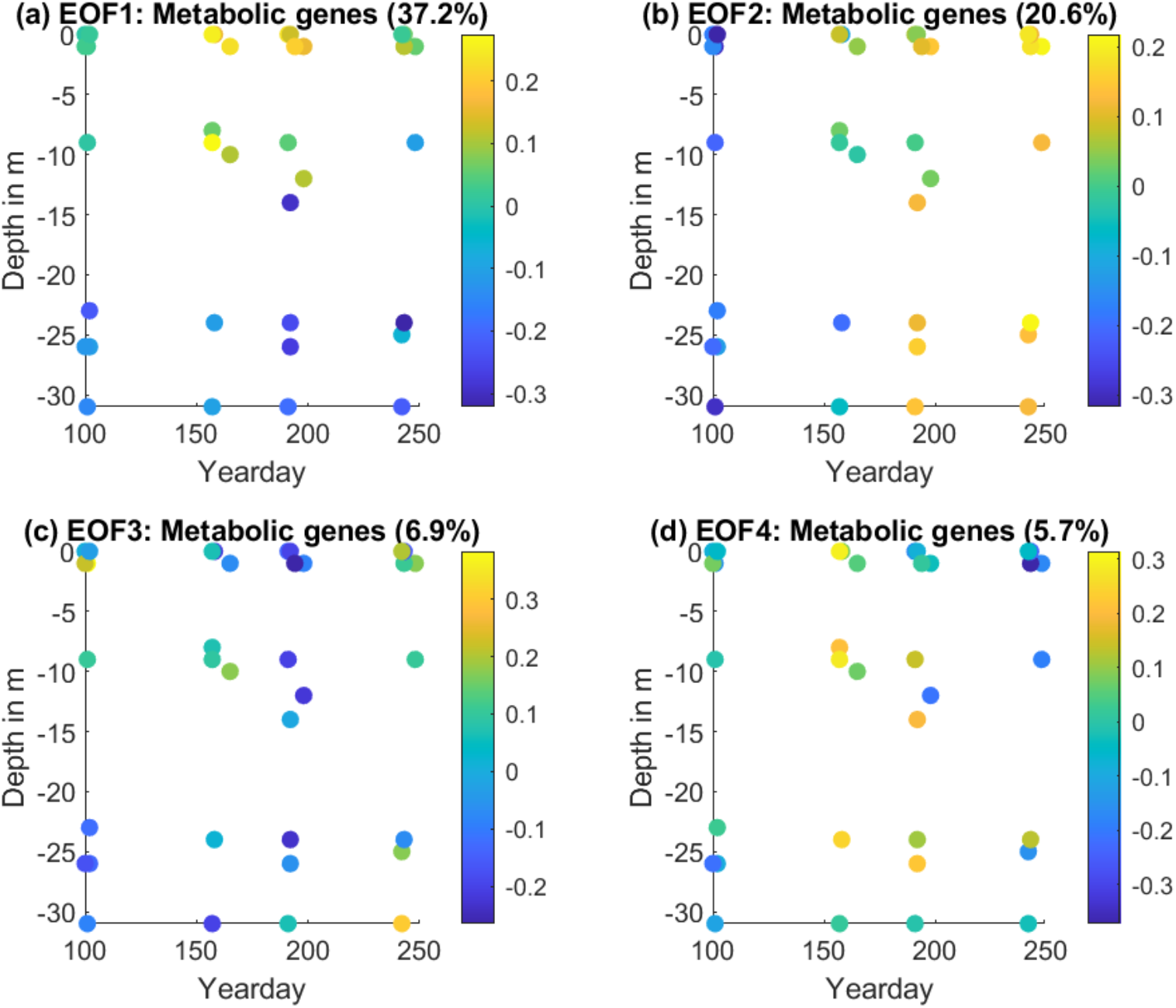
Leading patterns of variation with depth and time shown by the top four empirical orthogonal functions (EOFs 1-4) computed using the metabolic genes (excluding housekeeping genes) in Table 1. Colors represent EOF values and the percent of the variation explained by each EOF is indicated in parentheses in the title. Note that all stations are plotted together for each date.

The first EOF (Fig. 2a) is largely driven by differences in gene composition between surface and deep waters, with stronger stratification later in the summer than in the late spring. This pattern captures 37.2% of the variability amongst the 76 metabolic genes listed in Table 1. The second EOF, accounting for 20.9% of the variance, is largely driven by changes to gene relative abundance from spring to summer (Fig. 2b), contrasting late August with mid-April, with little dependence on depth. The remaining EOFs show a big drop off in variance explained, with the third and fourth EOFs explaining only 6.9% and 5.7%, respectively. The third EOF (Fig. 2c) contrasts genes with higher abundance in shallow waters in April and June and throughout the water column in August with those that are highest in the rest of the samples. The fourth (Fig. 2d) contrasts genes that are highest in April and August with those that are highest in June and July.

To determine which genes and environmental factors are associated with the largest changes in gene composition, we correlated the EOFs with relative gene abundances, looking for the highest positive or lowest negative correlations. EOF1 had the highest positive correlation with photosynthetic genes, including genes from photosystems I and II and photosynthetic electron transport, and largest negative correlations with genes associated with nitrification (*hao* and *amoC*, although not including *amoA* and *amoB* genes) and denitrification (*napA, norBC, nirKS, nosZ*), dissimilatory nitrate reduction to ammonia (*napA, nrfA*), and sulfur cycling (*dsrA, dsrB*). The correlation between EOF1 and housekeeping genes is relatively low (0.32) suggesting that the variation is not strongly driven by changes in gene copies per genome, but by the composition of the community. EOF1 seems to be driven largely by photosynthesis-dominated microbial communities in surface waters and decomposition-dominated microbial communities seen throughout the spring and summer, however the contrast between surface and deep samples is much larger in June, August and September than it is in April.

The second EOF is dominated by genes whose abundance increases over the course of the summer, explaining more than half of the variance of some photosynthesis genes (*psaC, petE, petG, petH, psbF*) and the methane oxidation/nitrification gene *pmoA/amoA* (these cannot been distinguished in KEGG). It also has a relatively weak correlation with the housekeeping genes (0.30). EOF3 correlates negatively with the nitrogen fixation gene *nifH* and positively with *pmoB/amoB* and *pmoC/amoC*. Finally, a different set of photosynthesis genes were the genes with the highest correlated to EOF4, suggesting that these genes are enhanced in both the late summer near the surface and spring near the bottom. EOF3 is weakly anticorrelated while EOF4 is weakly positively correlated with the housekeeping genes.

The two leading EOFs of the metabolic genes have strong relationships with several environmental variables. EOF1 is strongly correlated with pH, moderately correlated with log chlorophyll and anticorrelated with phosphate, apparent oxygen utilization (AOU) and ammonia. This is consistent with positive values from photosynthesis-dominated functions and negative values associated with decomposition and/or release of reduced substrates from the sediments (Table 2). We used a stepwise linear regression model in which at each step the environmental variable with the highest correlation to the data or to the residuals is added to the regression. We find that a combination of pH, phosphate and temperature predicts EOF1, with a correlation coefficient of 0.87 (note that while temperature is not very well correlated with mode 1, with a correlation of 0.22, its correlation with the residuals once the relationships with pH and phosphate are removed is 0.45). The second EOF is strongly correlated with temperature (0.88) and less correlated with phosphate (0.43) and dissolved organic phosphorus (0.43), both of which accumulate over the summer. These three variables were also found to predict EOF2 with a correlation coefficient of 0.91. To first order, modes explaining 58% of the variability in key metabolically-relevant genes are predictable from bulk physical, chemical, and biological measured values.

**Table 2:**
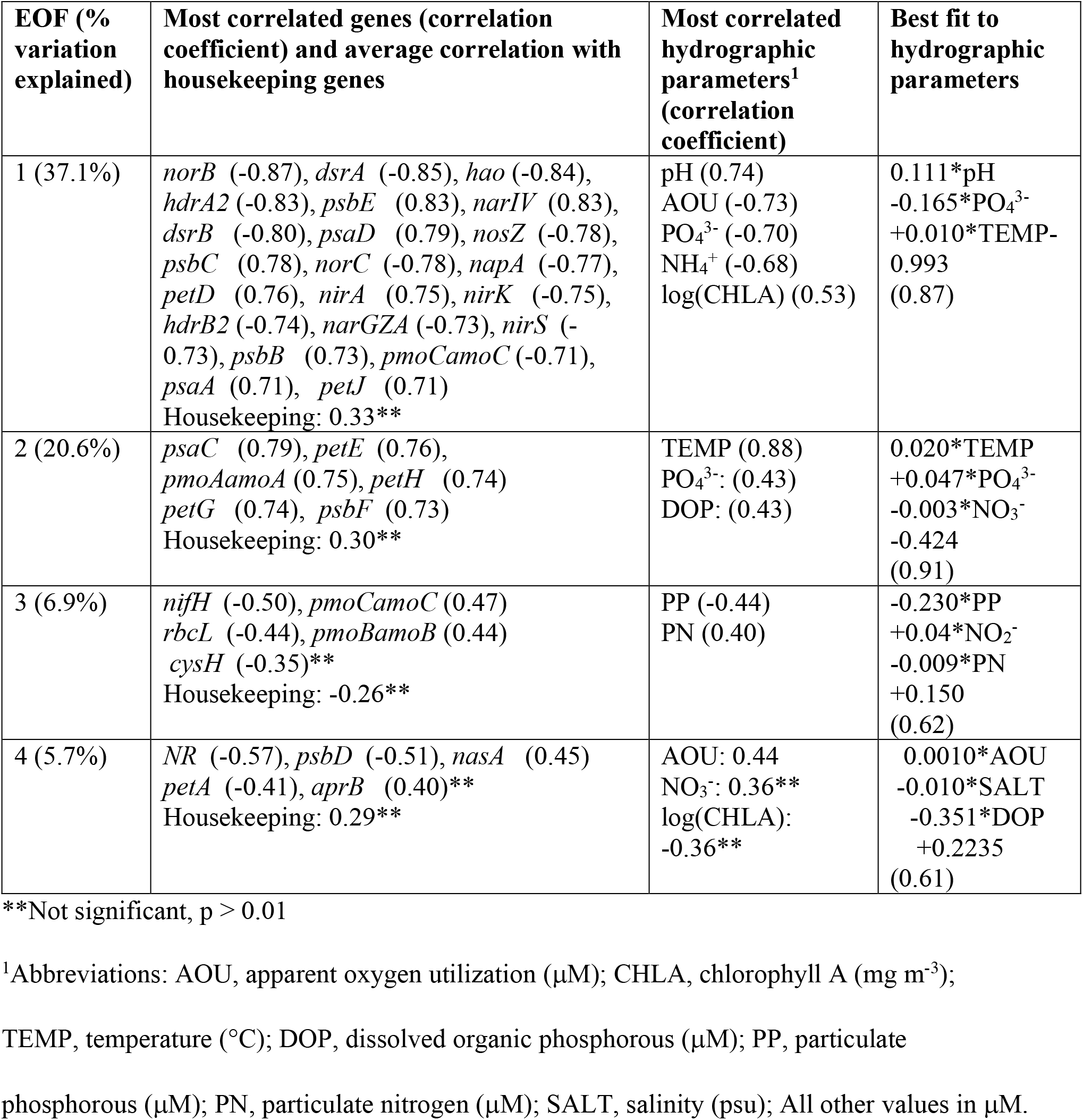
List of top EOFs (Column 1), leading gene correlates with each EOF (Column 2, showing all genes where the EOF either explains more than half the variance or the top five most correlated genes), most correlated hydrographic parameters (Column 3) and the best fit to three of the hydrographic parameters (Column 4).

| EOF (% variation explained) | Most correlated genes (correlation coefficient) and average correlation with housekeeping genes | Most correlated hydrographic parameters<sup>1</sup> (correlation coefficient) | Best fit to hydrographic parameters |
| --- | --- | --- | --- |
| 1 (37.1%) | *norB* (-0.87), *dsrA* (-0.85), *hao* (-0.84), *hdrA2* (-0.83), *psbE* (0.83), *narIV* (0.83), *dsrB* (-0.80), *psaD* (0.79), *nosZ* (-0.78), *psbC* (0.78), *norC* (-0.78), *napA* (-0.77), *petD* (0.76), *nirA* (0.75), *nirK* (-0.75), *hdrB2* (-0.74), *narGZA* (-0.73), *nirS* (-0.73), *psbB* (0.73), *pmoCamoC* (-0.71), *psaA* (0.71), *petJ* (0.71)<br>Housekeeping: 0.33** | pH (0.74)<br>AOU (-0.73)<br>PO<sub>4</sub><sup>3-</sup> (-0.70)<br>NH<sub>4</sub><sup>+</sup> (-0.68)<br>log(CHLA) (0.53) | 0.111*pH<br>-0.165*PO<sub>4</sub><sup>3-</sup><br>+0.010*TEMP-<br>0.993<br>(0.87) |
| 2 (20.6%) | *psaC* (0.79), *petE* (0.76), *pmoAamoA* (0.75), *petH* (0.74)<br>*petG* (0.74), *psbF* (0.73)<br>Housekeeping: 0.30** | TEMP (0.88)<br>PO<sub>4</sub><sup>3-</sup>: (0.43)<br>DOP: (0.43) | 0.020*TEMP<br>+0.047*PO<sub>4</sub><sup>3-</sup><br>-0.003*NO<sub>3</sub><sup>-</sup><br>-0.424<br>(0.91) |
| 3 (6.9%) | *nifH* (-0.50), *pmoCamoC* (0.47)<br>*rbcL* (-0.44), *pmoBamoB* (0.44)<br>*cysH* (-0.35)**<br>Housekeeping: -0.26** | PP (-0.44)<br>PN (0.40) | -0.230*PP<br>+0.04*NO<sub>2</sub><sup>-</sup><br>-0.009*PN<br>+0.150<br>(0.62) |
| 4 (5.7%) | *NR* (-0.57), *psbD* (-0.51), *nasA* (0.45)<br>*petA* (-0.41), *aprB* (0.40)**<br>Housekeeping: 0.29** | AOU: 0.44<br>NO<sub>3</sub><sup>-</sup>: 0.36**<br>log(CHLA):<br>-0.36** | 0.0010*AOU<br>-0.010*SALT<br>-0.351*DOP<br>+0.2235<br>(0.61) |

\*\*Not significant, $p > 0.01$
<sup>1</sup>Abbreviations: AOU, apparent oxygen utilization ( $\mu\text{M}$ ); CHLA, chlorophyll A ( $\text{mg m}^{-3}$ );
TEMP, temperature ( $^{\circ}\text{C}$ ); DOP, dissolved organic phosphorous ( $\mu\text{M}$ ); PP, particulate
phosphorous ( $\mu\text{M}$ ); PN, particulate nitrogen ( $\mu\text{M}$ ); SALT, salinity (psu); All other values in $\mu\text{M}$ .

It is also noteworthy that certain variables (salinity, phaeopigments, particulate phosphorus and nitrogen) are poorly correlated with both of the two leading modes of variability. The low correlation with salinity is particularly surprising as studies of taxonomy based on 16S rRNA gene sequencing (Arora-Williams et al. 2022; Wang et al. 2020) demonstrate salinity changes community structure, although previous work suggests taxonomic changes can often be uncoupled from functional changes (Louca et al. 2016b). In fact, the most significant correlation with salinity is only 0.36 and is associated with the 8^th^ mode of variability, which explains only 2.6% of the variability in metabolic genes. When looking at common genes, the most significant correlation with salinity (−0.69) is found in the fourth mode, which explains 5.8% of the variability.

Individual genes can also be predicted with skill from the environmental variables. Take for example *norB*, the gene which has the greatest correlation with EOF1 and which is associated with the production of nitrous oxide as a key step in denitrification. *NorB* can be predicted by the same variables that predict EOF1 with the equation:

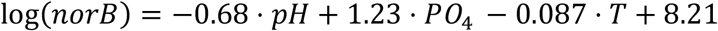

predicting the log frequency with a correlation coefficient of 0.86. This means that each increase in pH of 0.1 will lead to *norB* dropping by 7%, while an increase of phosphate of 0.1 mM will lead to an increase in *norB* of 13% and each increase of temperature by 1 °C will lead to a decrease in *norB* of 9%. While the maximum value of *norB* in the dataset is 265 times the minimum value, the standard deviation of the error gives a factor of 1.9 with the abundance at 30 of the 38 sites predicted to within a factor of 2. Similarly, the log abundance of *hao*, which plays a key role in nitrification, can be written as

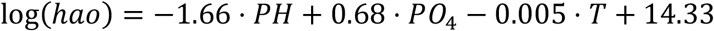

also with a correlation of 0.86. The common features then are that genes well predicted by EOF1 tend to have similar but opposite weightings of pH and PO_4_ and a weak dependence on temperature with the same sign as pH. The dependencies on PO_4_ and pH are not particularly surprising, suggesting that a gene associated with the production of nitrous oxide is found in waters that have high levels of decomposition products. However, the negative dependence on temperature is a surprise, and results in higher levels of *norB* during the April cruise, as we will discuss below.

We also compared how EOFs correspond to changes in the rate of key processes predicted by the model associated with carbon, nutrient, and sulfur cycling (Fig. S10). EOF1 correlates positively with photosynthesis rates in the model and most negatively with modeled ammonia oxidation and sulfate reduction rates. This result is partially supported by the gene abundance, where genes for photosynthesis (e.g. several *psb* and *pet* genes), ammonia oxidation (*hao* and *amoC*) and sulfate reduction (*dsrA*) are among the genes with the largest correlation coefficients with EOF1. Modeled denitrification rates do not show any significant correlation with EOF1, in contrast to the strong negative correlation of EOF1 with several key denitrification genes (e.g. *nar, nir, nor* and *nosZ* genes). The second and fourth EOFs show weaker non-significant (p > 0.01) positive correlations with sulfur metabolism. Abundance of genes associated with this process is low in April, highest in June and July and a little lower in August. Sulfur cycling genes were only identified as correlated to EOF4 (*aprB*), although also not significantly

### Patterns of genes associated with nitrification, denitrification and sulfur cycling compared to model predictions

A major advantage of shotgun metagenomics compared to our previous 16S rRNA gene analysis is the ability to investigate changes in the relative abundance of genes known to mediate biogeochemical transformations of interest. Instead of simply reporting on general patterns in genes or genomes across samples, we compared the relative abundance of key genes to predictions of the rates of the processes mediate by these genes generated from state-of-the-art models to guides our interpretation of these patterns. Changes in relative abundance of genes that correspond well with changes in corresponding process rates suggests that patterns are driven by the availability of energy to support the growth of these organisms (in the case of the subset of genes we chose), as has been previously suggested (Reed et al. 2014). Changes in relative abundance of specific genes that do not correspond well with corresponding rates suggest other factors are more important in determining the abundance of these genes or that there is an error in the model, both of which warrant further investigation.

#### Nitrification

Nitrification plays an important role in determining the fate of nitrogen after ammonification, contributes to the loss of oxygen from the water column, and facilitates denitrification. We focused on the first step in nitrification by examining four key genes involved in ammonia oxidation (*hao*, and *amoA, amoB*, and *amoC*) and comparing changes in relative abundance of these genes to changes in oxygen and modeled ammonia oxidation rates. The expectation is that these genes will be correlated with each other and with ammonia oxidation rates predicted by the model. For these expectations to hold, all genes would have to be present at a similar copy number in the genome of ammonia oxidizing microorganisms and the abundance of microorganisms across samples would need to be largely determined by the energy available from ammonia oxidation.

Seemingly in support of our assumptions, the log of *hao* gene abundance, which has been previously used as an index of nitrification (Arp et al. 2007), had a robust and significant positive correlation with log ammonia oxidation rates (0.712; Fig. 3b). However, log abundance of *amo* genes were less strongly correlated with log ammonia oxidation rates (*amoA*, 0.148; *amoB*, 0.257; *amoC*, 0.433). Additionally, while the log abundance of *amo* genes are relatively well correlated with each other (0.638-0.831), correlations with log abundance of *hao* are much weaker (*amoA*, 0.311; *amoB*, 0.432; *amoC*, 0.483). By digging into the dataset in more detail to better understand these discrepancies, we found evidence that several assumptions regarding the relationship of these genes with each other or the modeled ammonia oxidation rates were violated, likely contributing to some of the poor correlation values we observed, as described below.

**Figure 3.**
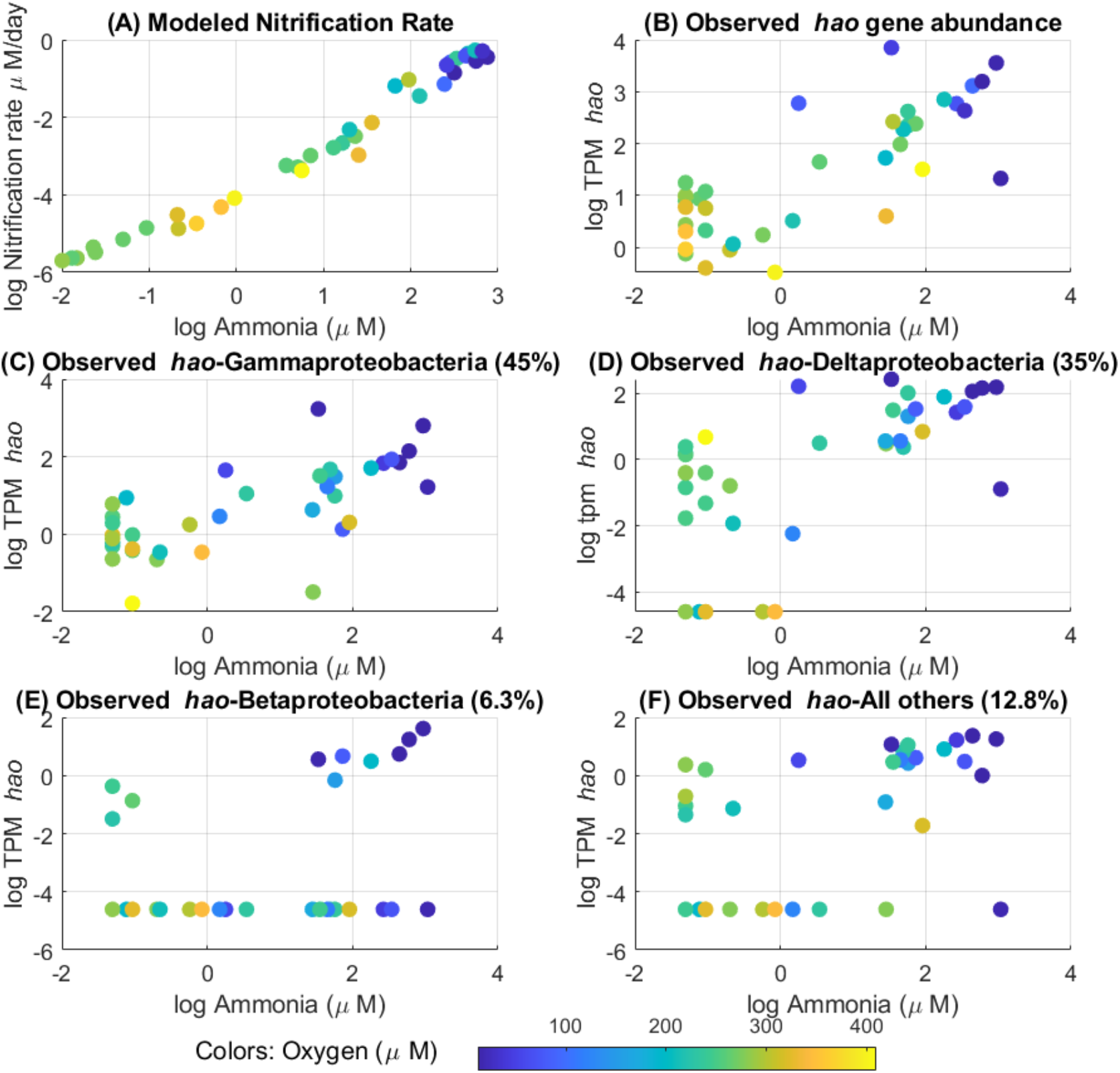
Relationship of (A) log modeled nitrification rate and (B) relative abundance of observed *hao* in log(TPM) related to log ammonia concentrations (μM, horizontal axis) and dissolved oxygen (μM, colors). (C-F) *Hao* relative gene abundance in log(TPM) separated by taxonomic groups; (C) Gammaproteobacteria, (D) Deltaproteobacteria, (E) Betaproteobacteria, and (F) all other taxonomic groups. Note that values at −4.6 represent zero values.

Many of the nitrification genes we examined are not likely to be present together in the genome of the same organism and many are not classified as known ammonia oxidizing taxa. While the taxonomic classification of *amoA* and *amoB* genes was dominated by the Archaea *Nitrosopumilus*, an ammonia oxidizer, the *hao* gene taxonomic classification was much more diverse and dominated by many Gammaproteobacteria taxa not known as ammonia oxidizers (Fig. 3c). 87.6% of the TPM signal from *amoA* across samples was classified as *Nitrosopumilus*, whereas Gammaproteobacteria accounted for 45.5% of abundance of *hao*. The dominant genera assigned to the *hao* group are Candidatus Thioglobus, Candidatus Riedellia and Helioglobus. It is not clear how these organisms use *hao*, but it is likely important to their response to high levels of ammonia in the system, as the log of gene abundance is strongly correlated with log ammonia concentrations (0.74). Archaea such as *Nitrosopumilus* do not have *hao* genes (Kozlowski et al. 2016; Walker et al. 2010), so are not expected to contribute to the signal of *hao*, although it is not clear which genes, if any, fulfill a similar role as *hao* in their genome. *AmoABC* genes are evolutionarily similar to methane monooxygenase genes (Holmes et al. 1995), which are not differentiated in KEGG, but methane oxidizing taxonomic groups made up only 0.9-4.0% of the total *amoABC* gene signal. Additionally, we searched for bacterial *amoABC* genes at the oxycline in another metagenomic dataset taken from multiple depths within the water column at CB3.3C (Arora-Williams et al. 2022), but did not find any *amoA* genes with a distribution that would suggest we are missing any taxonomic groups by probing the top and bottom of the water column (Fig. S11). Additionally, the model predicts ammonia oxidation to be relatively evenly distributed from below the thermocline to our sampled depths (Fig. S12), such that we are likely sampling in areas of the water column with the highest rates.

Another assumption, that these genes are found at the same copy number within the genome, is also likely to be violated, even for the Archaeal *amo* genes. Archaeal *amoC* is 3.5625-7.125 times more abundant than *amoA* and *amoB* (Fig. S13), especially in late summer. Archaea are known to have an additional copy of *amoC* (e.g. (Arp et al. 2007)), which could contribute to the discrepancy in the signal. However, these differences could also be related viral copies of *amoC* gene previously identified in another Chesapeake Bay metagenomic dataset (Ahlgren et al. 2019). One of the Archaeal *amoC* genes found in our dataset (NODE_18553_length_10238_cov_48.570657_9) is 100% identical to a previously identified viral *amoC* gene [Ga0129342_100020914 from (Ahlgren et al. 2019)] and is the most abundant *amoC* gene in the dataset. The viral *amoC* gene was also found to be expressed at a similar time of year in a previous metatranscriptomic dataset [(Hewson et al. 2014); Fig. S14]. Thus, the presence of *amoC* genes in viral auxiliary metabolic genes likely contributes to the differences in the abundance and correlation between *amo* gene subunits.

*Hao* and *amoC* were both correlated with the first EOF, thus contributing to some of the largest changes in gene abundance across the dataset. The strong relationship of *hao* with modeled ammonia oxidization rates is intriguing, especially because it was previously used as an index of ammonia oxidation. However, the differences between *hao* and *amo* genes as outlined above suggests that a better characterization of the microbial community accomplishing nitrification in this system is needed.

#### Denitrification

Many genes involved in denitrification (e.g. *nor*, *nar, nir, nos*) are strongly correlated with the first metabolic EOF, suggesting that changes in denitrification capabilities represent some of the largest genetic changes in the dataset. Denitrification in the model is governed by the concentration of nitrite, oxygen, and by the organic matter remineralization rate, making it unclear what the emergent relationships between hydrography and rates will be. Looking across our 38 sites, we see a strong negative relationship between oxygen and denitrification, and less of a relationship with the nitrite concentration (Fig. 4a). Denitrification is a multistep process, with the final two steps involving nitric oxide reductase, encoded by two genes *norB*, and *norC*, and nitrous oxide reductase, encoded by the *nosZ* gene. These genes can be found in organisms carrying out either complete or partial denitrification (Canfield et al. 2005). Since these genes are associated with the final steps of denitrification and were highly correlated with the first metabolic EOF, we focused on the relationship between *norB*, *norC*, *nosZ* and observed oxygen values or modeled denitrification rates.

**Figure 4:**
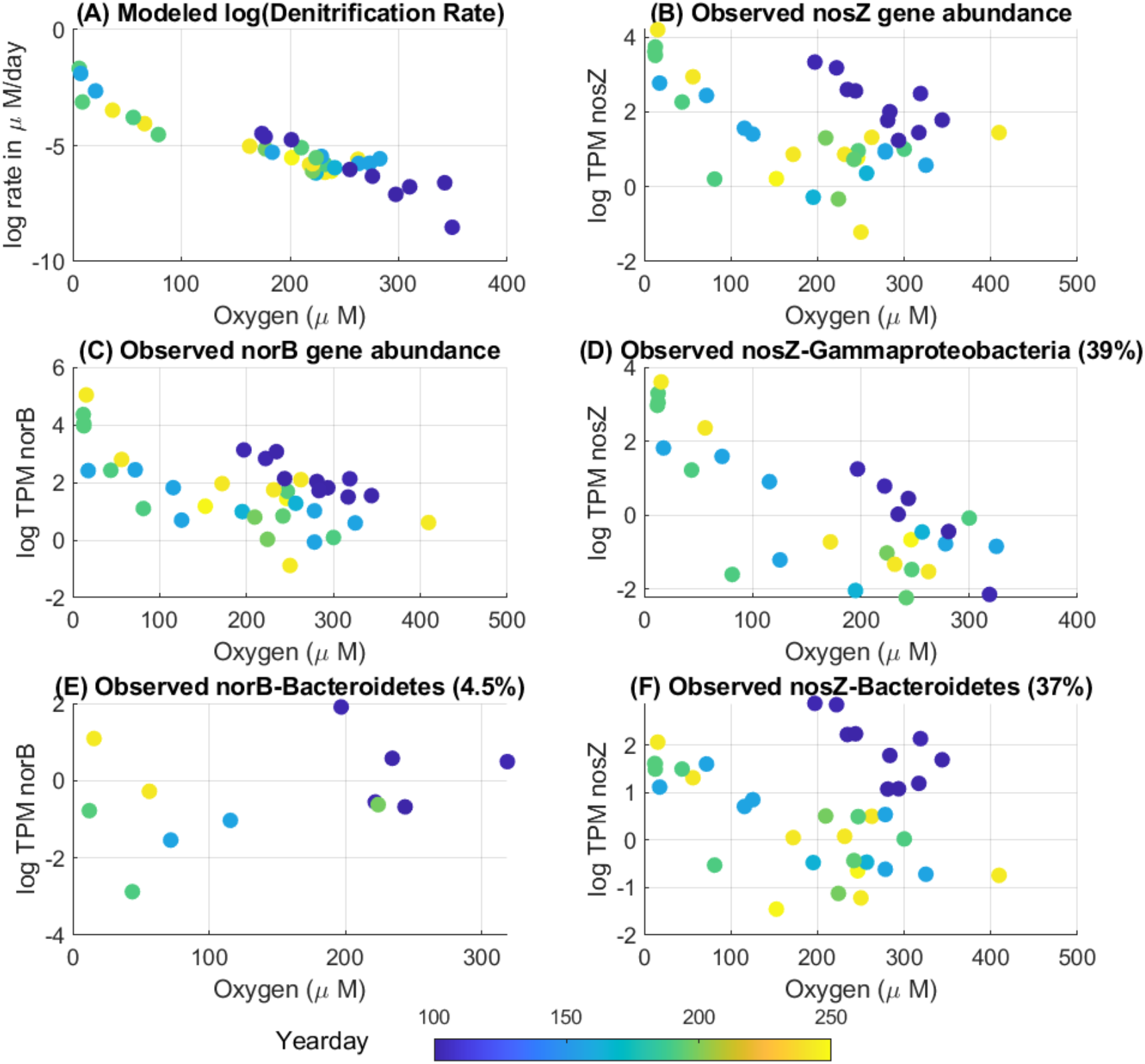
Modeled denitrification rates and key denitrification gene abundances related to oxygen concentration and time. Relationship of (A) log modeled denitrification rate and (B) observed *nosZ* gene frequency in log transcripts per million vs. dissolved oxygen (horizontal axis) and day of the year (colors). (C) Relationship of observed *norB* relative gene abundance in log(TPM) vs. dissolved oxygen (horizontal axis) and day of the year (colors). (D,F) Relationship of the subset of *nosZ* genes classified as (D) Gammaproteobacteria or (F) Bacteroidetes in log(TPM) vs. dissolved oxygen (horizontal axis) and day of the year (colors). (E) Relationship of the subset of *norB* genes classified as Bacteroidetes in log(TPM) vs. of dissolved oxygen (horizontal axis) and day of the year (colors).

We found that log abundances of denitrification genes are somewhat more highly correlated with each other than *amo* and *hao* genes, with *norB* well correlated both with *norC* (0.67) and *nosZ* well correlated with *nor* genes (*norB*, 0.88; *norC*, 0.68). By contrast, the correlation between log modeled denitrification and log abundance of *nosZ* (denitrification, 0.48; Fig. 4b) and *norB* (denitrification, 0.41; Fig. 4c) is not as strong as between *hao* and nitrification (0.712). Log denitrification decreases with increasing observed oxygen levels. While the highest relative abundances of *nosZ* and *norB* genes can be found at the lowest oxygen levels, *nosZ* shows relatively high abundance at intermediate levels of oxygen, largely associated with increased relative abundance in the spring (Fig. 4b, dark blue filled values). Log abundance of Gammaproteobacterial *nosZ* (Fig. 4d) has a stronger negative correlation with oxygen (−0.73) and a better positive correlation with log denitrification rates (0.69). However, *nosZ* associated with Bacteroidetes (Fig. 4e), mostly attributed to *Rhodothermus* and *Roseivirga*, increases to the highest levels at intermediate oxygen concentrations in the spring (Fig. 4f). Bacteroidetes dominate in the spring while Gammproteobacteria dominate in the summer. During the April cruise, 75.7% of the *nosZ* signal is associated with genes classified as Bacteroidetes. During the summer, Gammproteobacteria are 51.2% of the relative abundance *nosZ* and Bacteroidetes is only 21.8%. Thus while the overall correlation between the log of the modeled denitrification rate and log *nosZ* is relatively low (0.43), this can be broken down into *nosZ* associated with Bacteroidetes, which have a lower correlation with denitification rates (0.28), and *nosZ* associated with Gammaproteobacteria, which has a higher correlation (0.69). The results are similar for *norB* and *norC. NosZ* genes from this dataset had moderate expression in the spring in a previous dataset (Fig. S15), similar to the expression at certain points in the summer. Overall, our analysis suggests that while there is some agreement between oxygen or log denitrification rates with the log-abundance of denitrification genes, the model does not capture the relationship between denitrification and Bacteroidetes *nosZ*.

#### Sulfur cycling

Sulfate can be used as an electron acceptor during organic matter decomposition when oxygen is low, creating sulfide. Sulfide can then be oxidized, either biotically or abiotically, to oxidized forms of sulfur when oxygen or nitrate become available. While many Bay models only include abiotic sulfide oxidation (Da et al. 2018), we used a model that would allow biotic sulfide oxidation with both oxygen and nitrate. Key genes involved in both sulfate reduction and sulfide oxidation are dissimilatory sulfite reductase subunits alpha (*dsrA*) and beta (*dsrB*), which are highly correlated across samples (0.81) and strongly correlated with EOF1. The *dsrA* and *dsrB* genes are relatively more abundant in bottom samples from all cruises (Fig. 5a and b). During the summer, the relative abundance of *dsrAB* drops near the surface while building to high values in the late summer at deeper stations. It is worth noting that even during the April, the average relative abundance of these genes (36.3 ± 11.2 for *dsrA* and 5.4 ± 26.9 for *dsrB*) is comparable to *hao* (20.9±15.1) although not as abundant as *nosZ* (173.8±5.1), suggesting they may play an important biogeochemical role in relatively well oxygenated waters in the spring. As with *nosZ*, the taxonomic distribution is very different for April sites than for the rest of the season (Fig. S16). For example, in samples from all months except April, the most common taxonomic classifications for *dsrA* are *Thioploca*, accounting for almost 30% of the total, *Thiohalobacter*, accounting for 16%, and another 15% associated with an unclassified bacteria identified as an molluscan endosymbiont. In April, *Thiohalobacter* is still 17% of the signal but the other three account for only about 8% of the signal, a fivefold drop.

**Figure 5:**
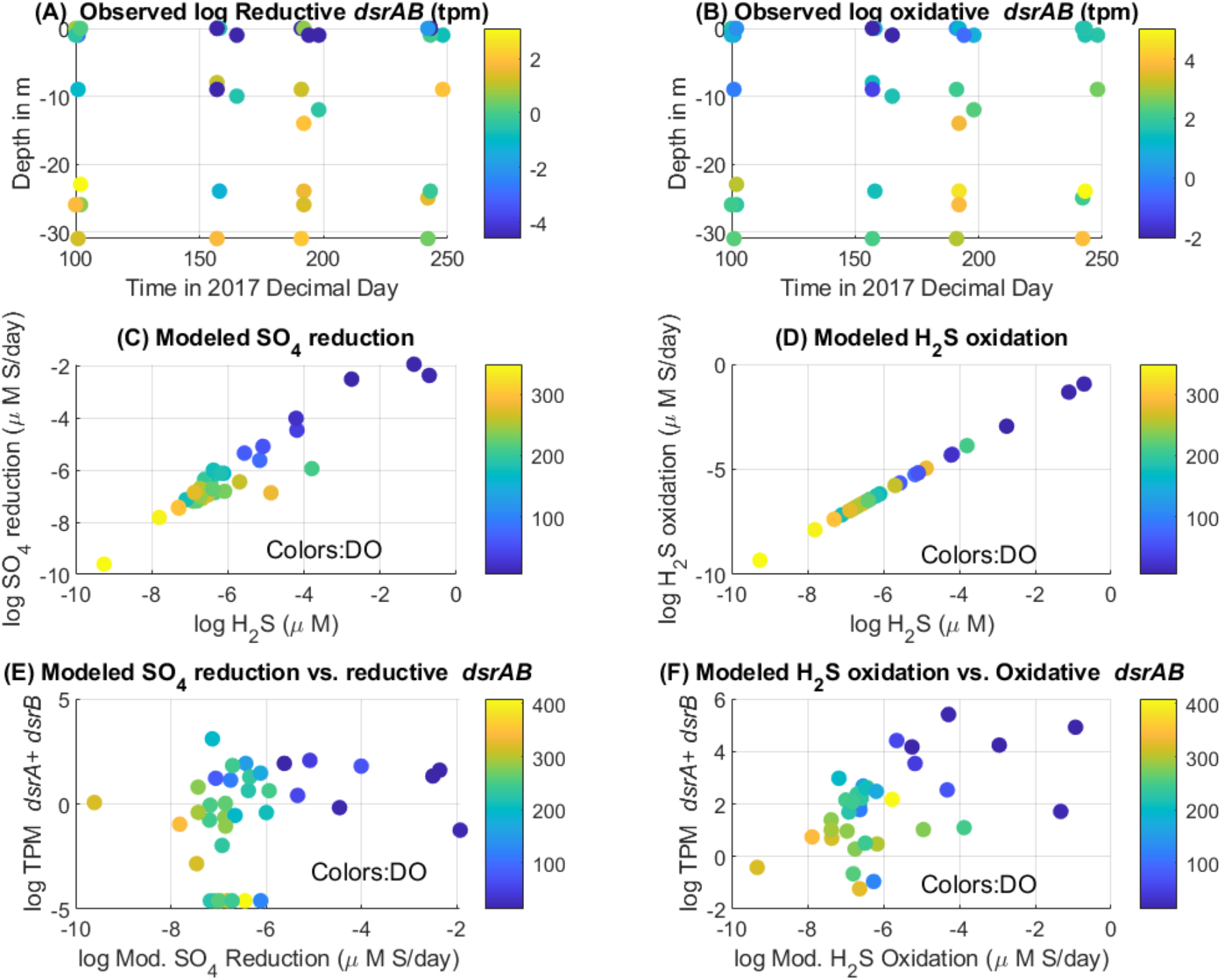
Distribution of genes in space and time (top row) and relationship of sulfate reduction or sulfur oxidation rates with gene abundance and predicted hydrogen sulfide concentration (middle and bottom rows). (A-B) Observed distribution of (A) *dsrA* and (B) *dsrB* with depth between April and August, with colors indicating the sum of both oxidative and reductive *dsr* types in log(TPM). Note that all stations are plotted together on each date. (C) Modeled sulfate reduction rates as a function of modeled H2S concentration, with colors showing modeled oxygen concentrations in μM. (D) Modeled sulfide oxidation (sulfide oxidation rates using nitrate, nitrite, and oxygen as oxidants are summed) as a function of modeled H2S concentration, with colors showing modeled oxygen concentrations in μM. (E) Modeled sulfate reduction rates vs. sum of observed reductive types of *dsrA* and *dsrB*. Colors indicate oxygen concentrations in μM. (F) Modeled sulfide oxidation rates vs. sum of observed oxidative types of *dsrA* and *dsrB*. Colors indicate oxygen concentrations in μM.

Our model simulates sulfate reduction in a similar manner to denitrification, setting it proportional to the remineralization of organic matter. Modeled sulfide oxidation is relatively tightly related to modelled H2S (Fig. 5c) indicative of a relatively short lifetime for sulfide. Because sulfide oxidation is simulated as a simple exponential decay, with the oxidants partitioned amongst oxygen, nitrate and nitrite, the modeled sulfide concentration is exactly proportional to the loss via sulfide oxidation (Fig. 5d). Comparing the modeled rates of sulfate reduction with reductive forms of *dsrAB* (Fig. 5e), there is a poor correlation between gene abundance and rates (0.26). However, comparing the modeled rates of sulfide oxidation with oxidative forms of *dsrAB* (Fig. 5f), we see a general consistency with sulfide oxidation, where modeled sulfide concentrations correlate well with log *dsrAB* relative gene abundances (0.51). When we try to analyze which form of *dsrA* is dominant, we find that that sulfide-oxidizing form accounts for about 85%, the sulfate-reducing form for about 7%, with the remainder being unassigned to either form. This is true throughout the season and across oxygen levels. The poor relationship between reductive *dsrAB* forms and sulfate reduction rates may be related to a low signal to noise ratio associated with low abundance genes. The overall picture is consistent with sulfur cycling playing a significant role in the Bay, as suggested by Arora-Williams et al. (Arora-Williams et al. 2022).

### Processes not included in the model

We now turn to processes which are not included in our model, but which might nonetheless play an important role in biogeochemical cycling.

#### Dissimilatory nitrate reduction to ammonia (DNRA)

Dissimilatory nitrate reduction to ammonia (DNRA) involves a set of nitrite reductases (*nirB, nirD, nrfA, nrfH*) that reduce nitrite to ammonia but are distinct from the nitrite reductases (*nirA*,NIT-6) involved in assimilatory nitrate reduction to ammonia. If we plot the four genes together against dissolved oxygen (Fig. 6a), a similar pattern to *nosZ* emerges, with non-April samples showing a strongly nonlinear increase as oxygen decreases but with April samples largely found above this curve (log-log correlation −0.61). The mean frequencies of these genes are surprisingly high, averaging 23 TPM across the whole dataset. Taken together, the strong relationship of DNRA genes with oxygen and their high relative abundance across the dataset suggest that the fate of nitrite within this system, specifically, whether it is nitrified, denitrified or returned to ammonia, requires further investigation.

**Figure 6:**
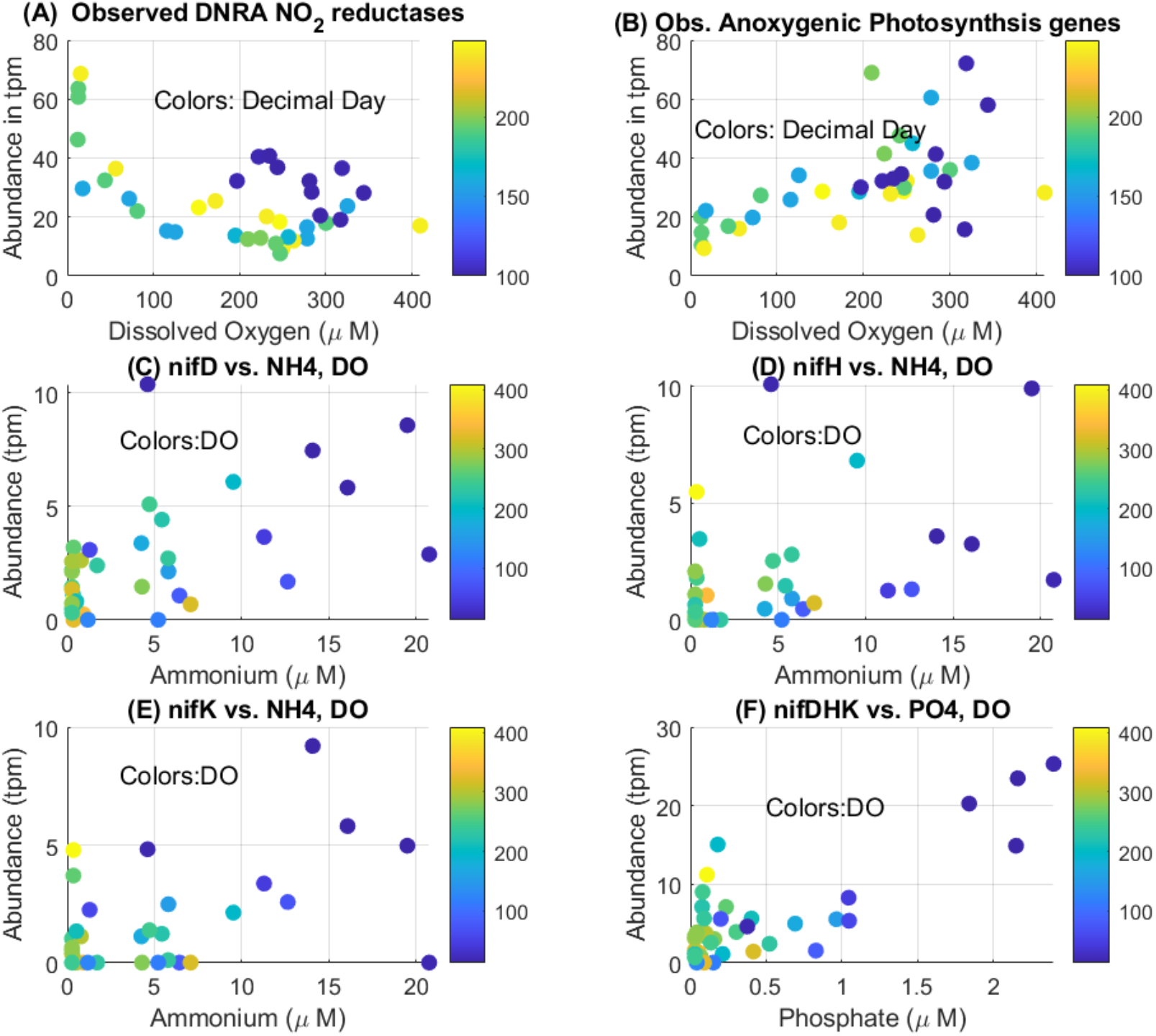
Genes associated with processes not currently resolved in the model. (A) Relative abundance of genes associated with DNRA (*nrfA+nrfH+nirB+nirD*) as a function of oxygen (horizontal axis) and day of the year (colors). (B) Relative abundance of anoxygenic photosynthesis genes (*pufL+pufM*) as a function of oxygen (horizontal axis) and day of the year (colors). (C-E) Relative abundance of nitrogen fixation genes (C) *nifD*, (D) *nifH*, (E) and *nifK* as a function of ammonium (horizontal axis) and oxygen (colors, in μM). (F) Sum of the three nitrogen fixation genes *nifD+nifH+nifK* as a function of phosphate (horizontal axis) and oxygen (colors, in μM).

#### Anoxygenic photosynthesis

Anoxygenic photosynthesis is a process by which bacteria use light to generate energy and oxidize reduced substrates such as sulfide, hydrogen, or iron. Previous work based on culture-dependent techniques suggested that a small population of anoxygenic photosynthetic Chlorobi would be sufficient to account for the anaerobic light-dependent sulfide oxidation in the Bay (Findlay et al. 2015). Two key photosystem II genes include the L (*pufL*) and M (*pufM*) subunits of the photosynthetic reaction center, which encode genes that can be appropriate markers for bacteriochlorophyll-mediated anoxygenic photosynthesis in organisms using the type II photosystem, such as Proteobacteria, Chloroflexi, and Gemmatimonadetes (Imhoff et al. 2018). These genes can be found across all stations in the dataset, with a positive correlation with oxygen (0.62 for *pufL* and 0.67 for *pufM*) and a maximum in April (Fig. 6b). We find a great deal of diversity of potential anoxygenic photosynthetic organisms, with 44 different species assigned to *pufL* and 50 assigned to *pufM*. The most common taxonomic assignment of *pufL* and *pufM* (accounting for about 15% of reads) is the species *Planktomarina*, a deeply branching Roseobacter species with the potential for aerobic anoxygenic photosynthesis (Giebel et al. 2019). The mean abundance of *pufLM* genes is comparable to core photosynthetic genes such as *psaA*, although not as abundant as *psbA*, suggesting that light-dependent oxidation of reduced chemical species, such as sulfide, in the Bay could be mediated by a more diverse group of microorganisms than previously recognized.

#### Nitrogen fixation

I Nitrogen fixation is a key bacterially mediated process that can act to balance the loss of nitrogen by denitrification, especially in environments with relatively high concentrations of biologically available phosphorous. This process can be carried out by photosynthetic bacteria or other bacteria in surface waters but has also been found to occur in sediments even with relatively high level of ammonium (Bertics et al. 2013; Gier et al. 2016; Knapp 2012). Nitrogen fixation involves a number of nitrogenases which transform nitrogen gas to ammonia. While these metalloenzymes can incorporate iron or vanadium as a key component, the most common genes and the only types we found in the Bay are *nifD, nifH*, and *nifK* which code for molybdenum-iron containing enzymes. We note that *nifD* is highly anti-correlated with the first EOF, changing dramatically over the season, and anticorrelated with the oxygenic photosynthetic genes, suggesting it is not associated with oxygenic photosynthetic organisms. The reads of all three nitrogenase-coding genes show a weak-to moderate positive correlation with ammonium (0.57, 0.45 and 0.51 for *nifD, nifH* and *nifK* respectively) and negative relationships with oxygen. All three genes tend to be found in the deep waters later in the season where there is a buildup of nutrients and pH is low. When we correlate the frequency of the three genes together against PO_4_, we get an even better relationship (Fig. 6f) with a correlation of 0.80. Nitrogen fixing genes were assigned to a diverse set of microorganisms, but sulfur-metabolizing bacteria were commonly found as some of the most abundant, including *Desulfomonas* (11.3%), *Sedimenticola* (6.0%), *Desulfobulbus* (4.3%) and *Desulfobacterium* (3.8%). It should be noted that nitrogen fixation is typically not included in many Bay models and was not encoded our new model.

## Discussion

Our metagenomic analysis of the Chesapeake Bay, a key estuarine ecosystem, has revealed that pH, PO_4_, and temperature can be used to explain the largest changes in relative abundance of key genes in space and time. By comparing how the relative abundance of key genes changes in relation to other genes in the same pathway and model-predicted rates for processes those genes are involved in, we were able to highlight areas of general agreement and point out discrepancies. There was general agreement between the log abundance of *hao*, a gene previously used as an index for nitrification, and the log of predicted nitrification rates in the model. Modeled denitrification rates were well correlated with the log relative abundance of non-Bacteroidetes *norBC* and *nosZ* genes and modeled sulfide oxidation rates were well correlated with oxidative types of *dsrAB*. Some of the largest discrepancies that could have substantial implications for biogeochemistry include the increase in relative abundance and expression of genes involved in 1.) anaerobic metabolism in well oxygenated waters in the spring, 2.) increase in relative abundance and expression of genes involved nitrogen fixation in high phosphate, low oxygen bottom waters, and 3.) discrepancies between *amo* genes, *hao*, and predicted nitrification rates. Although the cause and implication of these discrepancies cannot be determined through metagenomic analysis alone, these results highlight areas and processes for which our expectations about what is driving changes in gene abundance is not supported by our observations.

Our analysis advances the comparison of relative gene abundances and biogeochemical models of predicted rates detailing both similarities and discrepancies. We found that sub-setting gene abundances according to taxonomic classification in some cases could improve the relationship with modeled processes. This was also seen previously with relationships to measured nitrification potential, where ammonia-oxidizing bacteria were significantly correlated with nitrification potential, while ammonia-oxidizing archaea were not (Tatti et al. 2022). Certain taxonomic groups, like Bacteroidetes, could have a substantially different approach, such as being attached to or found within particles (Bianchi et al. 2018). Understanding the factors that influence the abundance of anomalous taxonomic groups could reveal unique approaches or responses to environmental factors that are not currently accounted for in models, such as microenvironments within particles. There has been a recognized need in ecological models to encode different phytoplankton groups separately [e.g. (Follows et al. 2007)], even though it has been met with challenges (Shimoda and Arhonditsis 2016). It could be important to consider encoding different taxonomic groups for other processes besides photosynthesis. Additionally, viral auxiliary metabolic genes could contribute to discrepancies between the abundances of genes in the same pathway or different subunits of the same enzyme (i.e. *amoABC* genes). Any effort to derive biogeochemical rates from gene copy numbers, similar to previous attempts (Reed et al. 2014), may need to consider the impact of viral auxiliary metabolic genes on this calculation.

Several caveats to our approach should be noted. Previous work reported changes in the absolute abundance of genes (Reed et al. 2014), rather than relative abundances reported here. This could become important if the number of cells changes dramatically between samples, or the average genome size changes. The low variation in single-copy housekeeping genes suggest a similar average genome size, while cell concentrations have been previously observed to vary less than an order of magnitude over time at one site in the Bay (Crump et al. 2007). Applying quantitative metagenomics with spiked internal standards has the potential to provide both high-throughput gene information and quantification to improve this comparison (Crossette et al. 2021). Another caveat of our approach is that the observed changes in DNA copy number of key genes does not take into account the activity of the respective enzymes and may include dead or inactive cells. Observations of expression of these genes in a previous metatranscriptomic dataset (Hewson et al. 2014) suggests the observed genes are actively expressed under similar conditions as our study, although even expression can diverge substantially from protein translation (Waldbauer et al. 2012) and may not reflect active processes (Rocca et al. 2015).

Our approach has also revealed discrepancies between the model and our observations that require additional work to better understand, but which could lead to exciting discoveries. The strong correlation between ammonia concentrations and *hao* genes is difficult to explain, especially since they were mostly associated with non-ammonia oxidizing taxonomic groups. One possibility is that the dominant ammonia oxidizing bacterial species have not been discovered, causing these *hao* genes to be classified as non-ammonia oxidizing taxa. If so, these organisms would need to have extremely divergent *amoABC* genes, since classified bacterial *amo* genes were much less abundant than *hao* genes. *Hao*-like genes in non-ammonia oxidizing taxa have been identified previously (Bergmann et al. 2005) and it is possible that these organisms may be using this multi-heme protein for its “off-label” electron storage capabilities, as previously suggested (Bewley et al. 2013). Additionally, the high abundance of *hao* in the Bay could indicate high levels of hydroxylamine, which may have gone undetected previously given that hydroxylamine has been relatively understudied (Soler-Jofra et al. 2021).

Both DNRA and denitrification displayed an overabundance of genes in well oxygenated waters in the spring compared to model predictions. Previous work in the Chesapeake Bay revealed the expression of genes related to the reduction of nitrate, iron, manganese, and sulfate during respiration occurred concurrently, rather in sequence as might be expected based on thermodynamic considerations (Eggleston et al. 2015; Hewson et al. 2014). This braiding of, rather than distinct transitions between, alternative electron acceptor processes was seen to occur within another dynamic and heterogeneous environment as well (Chen et al. 2017). Microaggregates creating anoxic microenvironments where many of the oxygen-sensitive processes could occur was thought to support the braiding of processes found in experimental mesocosms (Chen et al. 2017) and could also be an important source of denitrification in oxic waters (Liu et al. 2013). Thus, microenvironments on particles may explain the larger than expected gene abundances of anaerobic respiration processes in oxygenated water in the spring in our analysis.

We also observed an increase in the abundance of genes associated with nitrogen fixation deep in the water column in areas with high amounts of ammonia. Because nitrogen fixation is a more energetically costly pathway to assimilate nitrogen needed for anabolic processes compared to ammonia, this process was found to be inhibited by high concentrations of ammonia in the environment, as it is in model diazotrophs such as *Azotobacter* and *Trichodesmium* [e.g. (Hartmann et al. 1986; Holl and Montoya 2005; Mulholland et al. 2001)]. However, substantial amounts of nitrogen fixation have been observed at ammonia concentrations of as much as 200 μm ammonia [see (Knapp 2012) for a complete description of studies]. Additionally, microorganisms such as *Trichodesmium* have been observed to overcome nitrate inhibition in the presence of a high concentration of phosphorous (Knapp 2012; Knapp et al. 2012). The abundance of nitrogen fixation genes deep in the water column and increasing with higher concentrations of phosphate suggest that nitrogen fixation should be investigated as an additional nitrogen input into this system where nitrogen is carefully scrutinized.

Developing a more complex set of microbial processes in models based on these observations, including anaerobic metabolism, potentially on particles, in oxic waters and nitrogen fixation in non-photosynthetic bacteria could help resolve a number of challenges. The modeled total nitrogen to total phosphorous ratio in the spring is much higher than Redfield, and comparable to the 36-40:1 ratio seen in rivers entering the Bay (Moyer and Blomquist 2022). However, over the course of the summer this ratio drops to close to the Redfield ratio of 16:1, with total nitrogen dropping even as phosphate rises, a clear signal of denitrification. Permitting more of this denitrification to occur in the spring would alleviate the model’s tendency to overpredict nitrite (Fig. S1i). In addition, allowing nitrogen fixation during the summer in concert with denitrification would allow oxygen to be drawn down without the requirement that ammonia builds up to excessively high levels, something that is a general problem with the models studied in Jin et al. (Jin et al. 2022). Additionally, the implications of a previously under-appreciated source of nitrogen to the Bay through fixation suggests phosphate management may be the key to improving water quality within the Bay. Given our current understanding, the fact that there is excess nitrogen in the Bay through much of the year leads to a natural focus on curbing nitrogen inputs to the Bay. But if declines in nitrogen driven by curbing inputs would be compensated by more input from nitrogen fixers in high-phosphate waters, limiting phosphorus could be more beneficial to Bay water quality.

In summary, by analyzing metagenomic data collected from spring to fall at multiple sites within the Chesapeake Bay, we were able to determine the most important changes to gene abundances within this ecosystem, identify environmental factors associated with these changes, and identify similarities and discrepancies between modeled processes and observed gene abundances. A major mode of variation in key microbial genes involved energy conservation and metabolism was between the lower and upper water column in the summer, with intermediate abundances of these genes throughout the water column in the spring. pH, phosphate and temperature could be used to capture 75% and 82%, respectively, of the variation in the top two modes (EOF1,2) of variation in relative gene abundance. The relative gene abundance of a certain genes (e.g. *hao, dsrA*, Gammaproteobacterial *nosZ*) was relatively well correlated with the model prediction of the associated processes, although the correlation was worse for other genes in the same pathway (e.g. *amoABC*) or associated with other taxonomic groups (e.g. Bacteroidetes *nosZ*). Deviations between gene abundance observations and model predictions identified the spring as a time where gene abundances were higher than expected given the model predictions for denitrification and sulfur cycling. Additionally, nitrogen fixation genes were more abundant in the deep, low oxygen waters, which is not currently in Bay models. These observations indicate where research should be targeted in the future to better understand nutrient cycling and factors influencing oxygen dynamics and provides insight in the factors that influence the relationship between gene abundances and corresponding modeled processes.

## Acknowledgements

Maryland Department of Natural Resources and Old Dominion Water Quality Laboratory made the hydrographic measurements and generously provided water samples collected during their Chesapeake Bay water quality monitoring program. Funding for this research was provided by the Johns Hopkins University, Institute for Data Intensive Engineering and Science (IDIES) under their seed funding program to S.P. and A.G. Support for sequencing was provided by the NSF Integrative Graduate Education and Research Traineeship (Water, Climate, and Health: Grant 1069213). S.P. and K.A-W received support from Maryland Sea Grant under award 131160 from the National Oceanic and Atmospheric Administration, U.S. Department of Commerce. The authors declare no conflicts of interest.

