## Supplemental Information for "Major trends and environmental correlates of spatiotemporal shifts in the distribution of genes compared to a biogeochemical model simulation in the Chesapeake Bay"

<u>Environmental variables change substantially over the sampling period</u>

There were significant changes in the measured environmental variables over the course of the spring and summer of 2017 (Figure S3). Water temperature is relatively cold and homogeneous across all depths and stations in the spring (Fig. S3a). As summer progresses, the water warms and becomes stratified into the summer months, but thermal stratification breaks down towards the end of the summer. Salinity exhibits a general decline from south to north but does not have a strong trend with depth or time (Fig. S3b). Chlorophyll (Fig. S3c) shows higher values at greater depths during the spring followed by a summertime bloom in surface waters and low values at depth. The stratification of the water column is associated with a decline in oxygen (Fig S3d), a buildup of phosphate and ammonium (Figs. S3e,f), and reduction in pH (Fig. S3g). In contrast with phosphate and ammonium, nitrate (Fig. S3h) does not accumulate in deep waters during the summer, reflecting the impact of denitrification. However, nitrite (Fig. S3i) starts off relatively low and builds up over the summer in deep waters in a manner similar to phosphate. Runoff values close to the climatological average during 2017 resulted in a relatively typical year in terms of the volume of hypoxic waters across the Bay (https://www.vims.edu/research/topics/dead_zones/forecasts/report_card/index.php ).

<u>Model captures many of the observed chemical changes, with largest errors simulating nutrient concentrations</u>

The model simulation with additional microbial nitrogen and sulfur cycling processes captures many of the major changes in the observed environmental variables. The changes in temperature and pH are well simulated by the model across the 38 samples, with a coefficient of determination (1-root mean squared error/standard deviation of the observations) exceeding 0.99 with relatively little dependence on season (Fig. S1). Salinity and oxygen can also be simulated relatively well, with some low bias to salinity and high biases in oxygen primarily in the summer. However, more problems arise in the simulation of nutrients, a common problem in models of the Bay (Irby et al., 2016). Nitrate has a high bias during the spring, and ammonium a high bias during the summer. In the Jin et al. (2022, submitted) model, we focused on improving simulations of oxygen, nitrate and ammonia, as these had been included in the previous version of the model. While all three of these fields improve in the new model, the improvement in nitrate results from nitrite (not previously simulated in the code) building up to unrealistic levels during the spring. This raises the question of whether the new model, while including processes such as sulfide oxidation with denitrification, sets the threshold levels for these too low. Additionally, more work needs to be done optimizing the initial phosphate field in the Bay, though phosphate is generally not a limiting nutrient in our code.

EOF analysis of the most abundant genes

We conducted EOF analysis on a larger subset of 8714 genes that appear in at least 30 of the 38 stations to identify patterns and drivers of change across all abundant genes to compare to just the metabolic gene results. The first mode of the metabolic genes correlates best with the third mode of the common genes (0.84), which explains only 7.5% of the variation of these genes. By contrast, the first mode of variation in the common genes (which explains 16% of the total variation) correlates best with the second (0.58) and fourth modes (0.55) of the metabolic genes and is strongly correlated with housekeeping genes (0.78) with a slope of 0.59. This mode of variation may be driven more by changes to genome size (e.g. more eukaryotic microorganisms with larger genomes) that changes in composition.

**Supplemental Tables**

**Table S1.** List of stations, locations, dates, and key environmental conditions.

| Template | Station Name | Month Year | DO (mg/L) | CHLA (µg/L) | NH4F (mg/L) | NO3F (mg/L) | pH[1] |
|---|---|---|---|---|---|---|---|
| 7.10.17 CB6.2 Bottom | CB62 | 07 17 | 2.6 | 2.848 | 0.0901 | 0.0026 | 7.56 |
| 4.10.17 CB6.2 Surface | CB62 | 04 17 | 10.14 | 4.806 | 0.0038 | 0.0001 | 8.3 |
| 6.15.17 CB6.2 Bottom | CB62 | 06 17 | 4.01 | 2.848 | 0.0166 | 0.0012 | 7.63 |
| 4.10.17 CB6.2 Bottom | CB62 | 04 17 | 9.07 | 7.2624 | 0.0038 | 0.0001 | 8.19 |
| 7.10.17 CB6.2 Surface | CB62 | 07 17 | 7.74 | 8.544 | 0.0038 | 0.0001 | 8.2 |
| 6.15.17 CB6.2 Surface | CB62 | 06 17 | 9.12 | 6.6216 | 0.0038 | 0 | 8.31 |
| 8.28.17 CB6.2 Surface | CB62 | 08 17 | 7.88 | 5.6604 | 0.0038 | 0.0001 | 8.03 |
| 8.28.17 CB6.2 Bottom | CB62 | 08 17 | 4.88 | 2.403 | 0.0812 | 0.0049 | 7.73 |
| 6.5.17 CB7.3 Bottom | CB73 | 06 17 | 6.24 | 2.136 | 0.0046 | 0.0003 | 7.86 |
| 4.10.17 CB7.3 Bottom | CB73 | 04 17 | 8.99 | 2.5632 | 0.0038 | 0.0001 | 8.07 |
| 7.10.17 CB7.3 Surface | CB73 | 07 17 | 7.17 | 5.9808 | 0.0038 | 0.0001 | 8.18 |
| 7.10.17 CB7.3 Bottom | CB73 | 07 17 | 6.7 | 1.8156 | 0.0073 | -0.0001 | 7.95 |
| 4.10.17 CB7.3 Surface | CB73 | 04 17 | 9.39 | 2.136 | 0.0038 | 0.0001 | 8.08 |

| | | | | | | | |
|---|---|---|---|---|---|---|---|
| 6.5.17 CB7.3 Surface | CB73 | 06 17 | 8.21 | 3.4176 | 0.0038 | 0 | 8.15 |
| 8.28.17 CB7.3 Surface | CB73 | 08 17 | 8 | 4.9128 | 0.0038 | 0.0001 | 8.09 |
| 8.28.17 CB7.3 Bottom | CB73 | 08 17 | 6.6 | 4.3788 | 0.0167 | 0.0018 | 7.95 |
| 4.12.17 CB3.3C Bottom | CB33C | 04 17 | 6.3 | 6.23 | 0.13333333 | 0.08985 | 7.3 |
| 4.12.17 CB3.3C Surface | CB33C | 04 17 | 10.2 | 2.136 | 0.099 | 0.7823 | 7.7 |
| 6.6.17 CB3.3C Surface | CB33C | 06 17 | 8.9 | 18.583 | 0.06 | 0.1848 | 8.3 |
| 7.10.17 CB3.3C Surface | CB33C | 07 17 | 7.9 | 17.302 | 0.005 | 0.0008 | 8.2 |
| 8.31.17 CB3.3C Surface | CB33C | 08 17 | 13.1 | 37.594 | 0.005 | 0 | 8.7 |
| 7.10.17 CB3.3C Bottom | CB33C | 07 17 | 0.39 | 1.282 | 0.273 | 0.00285 | 7.2 |
| 6.6.17 CB3.3C Bottom | CB33C | 06 17 | 0.57 | 0.62 | 0.2905 | 0.02855 | 7.1 |
| 8.31.17 CB3.3C Bottom | CB33C | 08 17 | 0.5 | 1.068 | 0.0645 | 0.035 | 7.4 |
| 8.30.17 CB5.3 Surface | CB53 | 08 17 | 7.4 | 5.474 | 0.024 | 0.0174 | 8 |
| 8.28.17 CB5.3-1 Bottom | CB53 | 08 17 | 5.5 | 2.483 | 0.0595 | 0.0077 | 7.9 |
| 4.10.17 CB4.3C Bottom | CB43C | 04 17 | 7.1 | 9.612 | 0.076 | 0.0222 | 7.6 |
| 4.10.17 CB5.3-1 Bottom | CB53 | 04 17 | 7.8 | 8.544 | 0.066 | 0.0206 | 7.7 |

| | | | | | | | |
|---|---|---|---|---|---|---|---|
| 4.10.17 CB5.3 Surface | CB53 | 04 17 | 11 | 11.214 | 0.013 | 0.0192 | 8.2 |
| 6.6.17 CB5.3-1 Bottom | CB53 | 06 17 | 3.7 | 1.068 | 0.073 | 0.0171 | 7.5 |
| 6.6.17 CB5.3 Surface | CB53 | 06 17 | 8.9 | 10.833 | 0.007 | 0.0555 | 8.4 |
| 4.12.17 CB4.3C Surface | CB43C | 04 17 | 11.7 | 13.73 | 0.007 | 0.3342 | 8.1 |
| 7.10.17 CB4.2C Bottom | CB42C | 07 17 | 0.41 | 0.62 | 0.225 | 0.00105 | 7.3 |
| 7.10.17 CB4.3C Bottom | CB43C | 07 17 | 0.41 | 0.62 | 0.197 | 0.0022 | 7.3 |
| 8.28.17 CB4.4 Surface | CB44 | 08 17 | 8.4 | 9.612 | 0.005 | 0.0006 | 8.2 |
| 7.10.17 CB4.4 Surface | CB44 | 07 17 | 9.6 | 14.685 | 0.011 | 0.0008 | 8.4 |
| 8.28.17 CB4.4 Bottom | CB44 | 08 17 | 1.8 | 1.495 | 0.018 | 0.053 | 7.5 |
| 7.10.17 CB4.4 Bottom | CB44 | 07 17 | 1.4 | 0.641 | 0.158 | 0.0075 | 7.4 |
| 6.6.17 CB4.4 Surface | CB44 | 06 17 | 10.4 | 14.952 | 0.005 | 0.0368 | 8.6 |
| 6.6.17 CB4.4 Bottom | CB44 | 06 17 | 2.3 | 1.831 | 0.177 | 0.0175 | 7.3 |
| 4.10.2017 CB4.4 Bottom | CB44 | 04 17 | 7.5 | 7.476 | 0.081 | 0.0141 | 7.7 |
| 6.15.17 Bay Bridge 0mR1 | CB33C | 06 17 | NA | NA | NA | NA | NA |
| 6.15.17 Bay | CB33C | 06 17 | NA | NA | NA | NA | NA |

| | | | | | | | |
|---|---|---|---|---|---|---|---|
| Bridge 14m | | | | | | | |
| 6.15.17 Bay Bridge 8m | CB33C | 06 17 | NA | NA | NA | NA | NA |
| 4.19.18 Negative Control | NA | NA | NA | NA | NA | NA | NA |
| 5.15.18 Negative Control | NA | NA | NA | NA | NA | NA | NA |
| 5.18.18 Negative Control | NA | NA | NA | NA | NA | NA | NA |
| 5.8.18 Negative Control | NA | NA | NA | NA | NA | NA | NA |

[1] Full list of sample attributes can be found in the DataVerse website associated with this publication.

**Table S2.** Files and provenance associated with Hewson et al. 2014 metatranscriptomics dataset.

| SRA accession | Metadata based label | Files | Personal Communication | Dropped |
|---|---|---|---|---|
| SRR984835 | 5/17/10_Oxic_3m_1 | 1 | FALSE | FALSE |
| SRR984837 | 5/17/10_Oxic_3m_2 | 1 | FALSE | FALSE |
| SRR984833 | 5/17/10_Oxic_13m_2 | 1 | FALSE | FALSE |
| SRR980293 | 5/17/10_Oxic_13m_1 | 1 | FALSE | FALSE |
| SRR984839 | 6/7/10_Anox_16.5m_1 | 1 | FALSE | FALSE |
| SRR1011421 | 9/21/11_Oxic_17m_1 | 2 | FALSE | FALSE |
| SRR1011422 | 9/21/11_Oxic_17m_2 | 2 | FALSE | FALSE |
| SRR984891 | 6/7/10_Anox_16.5m_2 | 1 | FALSE | FALSE |
| SRR985022 | 7/11/10_Anox_17.5m_1 | 1 | FALSE | FALSE |
| SRR985023 | 7/11/10_Anox_17.5m_2 | 1 | FALSE | FALSE |
| SRR985024 | 7/11/10_Oxic_3m_1 | 1 | FALSE | FALSE |
| SRR985025 | 7/11/10_Oxic_3m_2 | 1 | FALSE | FALSE |
| SRR985026 | 8/5/10_Anox_22m_1 | 1 | FALSE | FALSE |
| SRR985027 | 8/5/10_Anox_22m_2 | 1 | FALSE | FALSE |
| SRR985028 | 8/30/10_Oxic_3m_1 | 1 | FALSE | FALSE |
| SRR985029 | 8/30/10_Oxic_3m_2 | 1 | FALSE | FALSE |
| SRR985030 | 8/30/10_Anox_20m_1 | 1 | FALSE | FALSE |
| SRR985031 | 8/30/10_Anox_20m_2 | 1 | FALSE | FALSE |
| SRR985301 | 10/18/10_Oxic_13m_1 | 1 | FALSE | FALSE |
| SRR985423 | 10/18/10_Oxic_13m_2 | 1 | FALSE | FALSE |
| SRR988005 | 7/08/11_Oxic_3m_1 | 2 | FALSE | TRUE |
| SRR988007 | 5/24/11_Anox_18m_1 | 2 | FALSE | FALSE |
| SRR988006 | 7/08/11_Oxic_3m_2 | 2 | FALSE | TRUE |
| SRR988008 | 4/18/11_Oxic_13.5m_1 | 2 | FALSE | TRUE |
| SRR988009 | 4/18/11_Oxic_13.5m_2 | 2 | FALSE | TRUE |
| SRR988014 | 5/24/11_Anox_18m_2 | 2 | FALSE | FALSE |
| SRR988024 | 6/14/11_Oxic_3m_2 | 1 | FALSE | FALSE |
| SRR988025 | 6/14/11_Anox_18m_1 | 2 | FALSE | FALSE |
| SRR988026 | 6/14/11_Anox_18m_2 | 2 | FALSE | FALSE |
| NA | 4/18/11_Oxic_13.5m_1 | 2 | TRUE | FALSE |
| NA | 4/18/11_Oxic_13.5m_2 | 2 | TRUE | FALSE |
| NA | 6/14/11_Oxic_3m_1 | 2 | TRUE | FALSE |
| NA | 7/8/11_Oxic_3m_1 | 1 | TRUE | FALSE |
| NA | 8_30_11_18m_18 | 1 | TRUE | TRUE |
| NA | 7/8/11_Oxic_3m_2 | 2 | TRUE | FALSE |
| NA | 8/30/11_Oxic_3m | 2 | TRUE | FALSE |
| NA | 8/30/11_Oxic_18m | 2 | TRUE | FALSE |
| NA | 8/8/11_Anox_18m_1 | 2 | TRUE | FALSE |
| NA | 8/8/11_Anox_18m_2 | 2 | TRUE | FALSE |

### Supplemental Figures

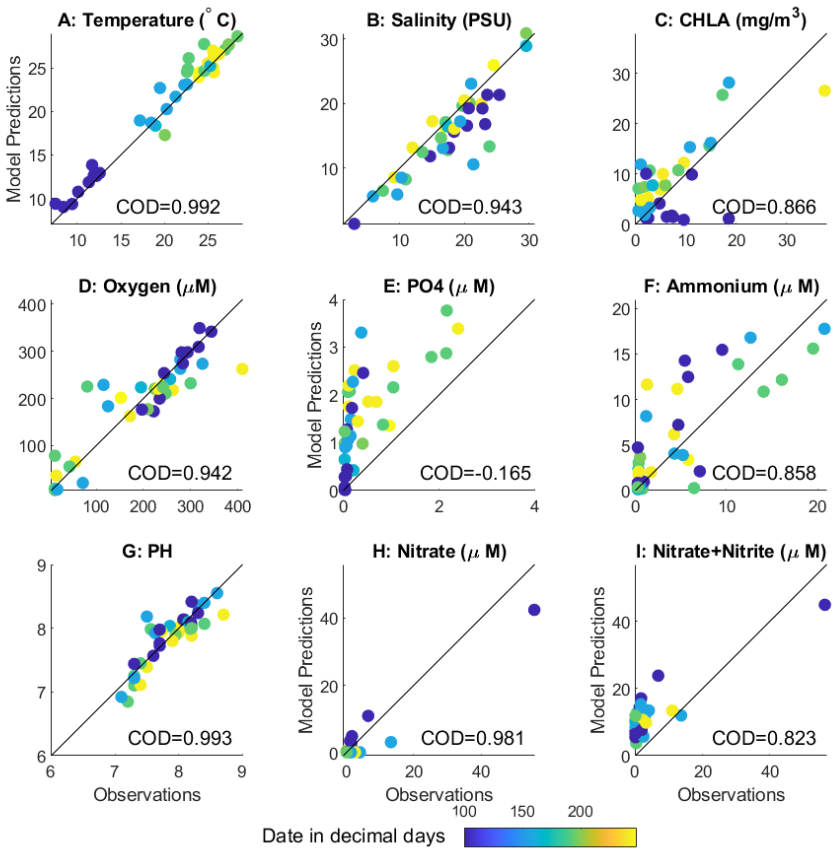


**Figure S1:** Comparison of observed (horizontal axis) and modeled (vertical axis) fields paralleling Figure 1. (A) Temperature (B) Salinity (C) Chlorophyll-a (D) Dissolved oxygen (E) Dissolved phosphate (F) Ammonium (G) pH (H) Nitrate and (H) Nitrate+Nitrite. Coefficient of determination is 1-mean square error/variance of observations. Line shows a 1:1 relationship.

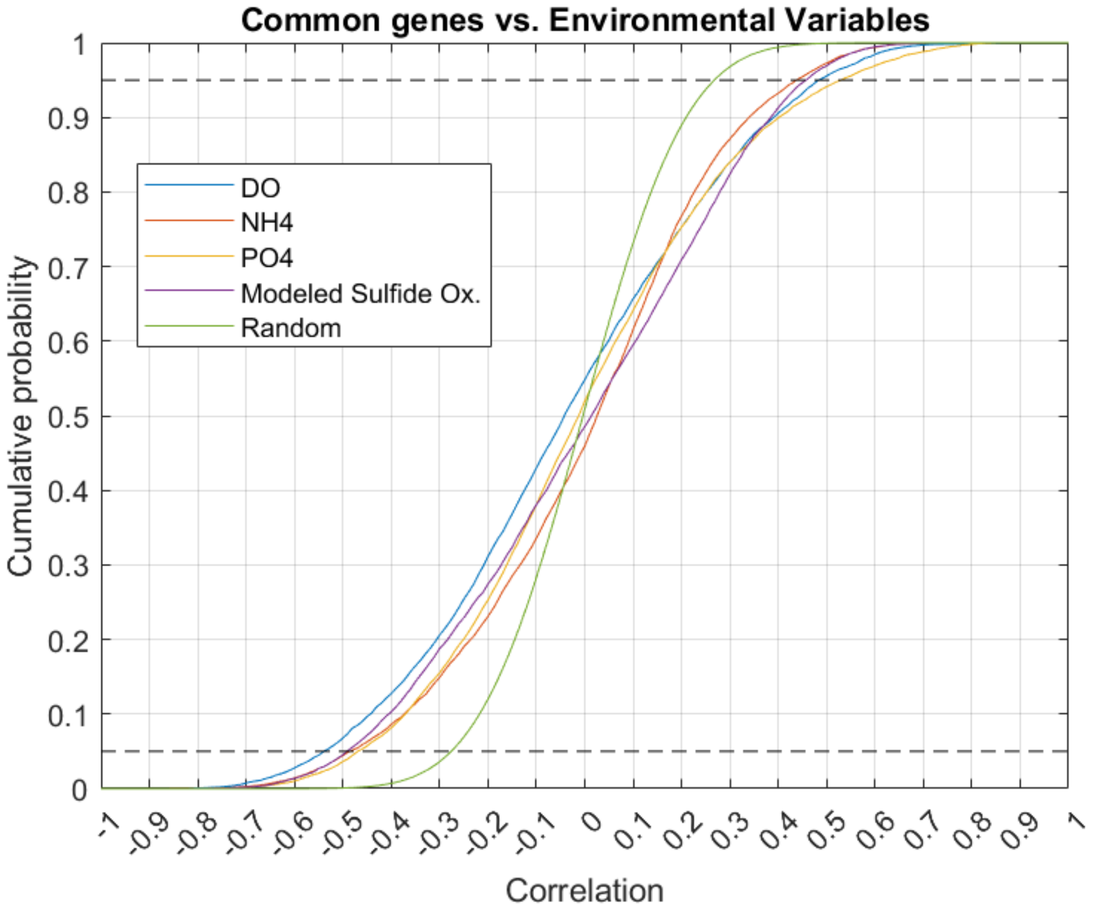


**Figure S2.** Cumulative probability curve of the correlation all 8,714 abundant genes with various observed and modeled values. Dotted lines represent top and bottom 5% of correlations. DO, dissolved oxygen; NH4, ammonium; PO4, phosphate; Modeled Sulfide Ox., Modeled sulfide oxidation rate; Random, a randomly generated set of datapoints.

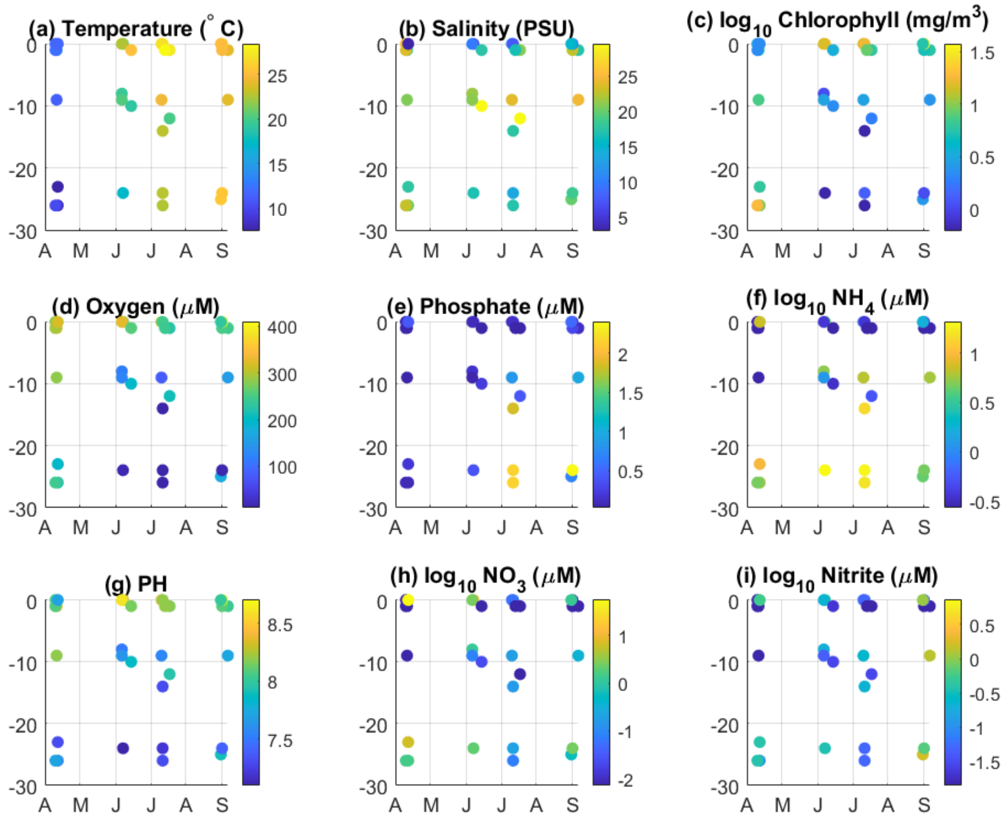


**Figure S3:** Summary of hydrographic measurements plotted against time (horizontal axis) and depth (vertical axis) for (a) temperature, (b) salinity, (c) log chlorophyll-a, (d) dissolved oxygen, (e) phosphate, (f) log ammonium, (g) pH, (h) log nitrate, and (I) log nitrite. Log scales are used for chlorophyll, nitrate, ammonium and nitrite to keep plot from being dominated by a few extreme values. Note, the spatial resolution between samples is not represented in this view, as all stations are plotted simultaneously for each date.

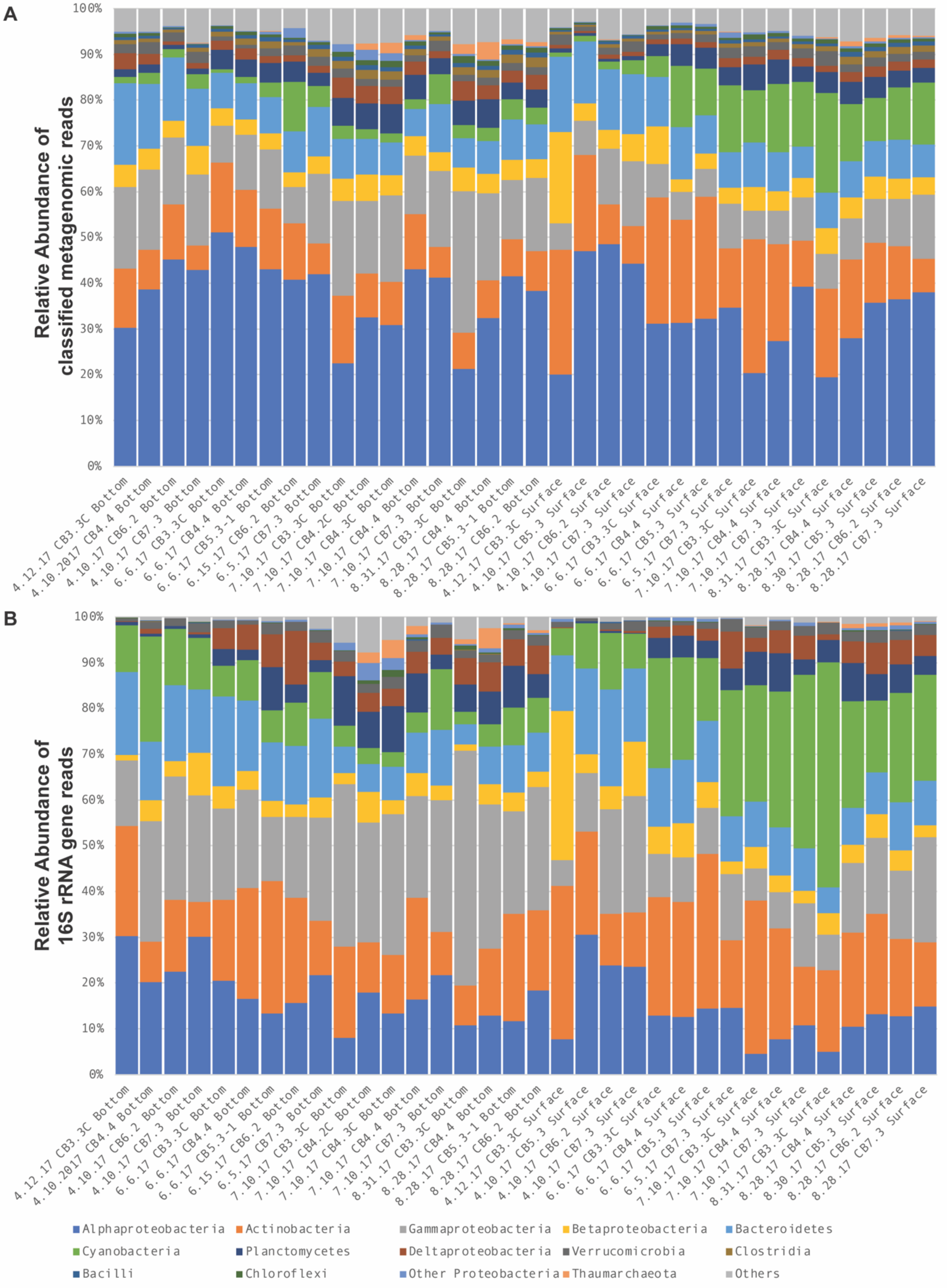
A
Relative Abundance of classified metagenomic reads
100%
90%
80%
70%
60%
50%
40%
30%
20%
10%
0%
4.12.17 CB3.3C Bottom
4.10.2017 CB4.4 Bottom
4.10.17 CB6.2 Bottom
4.10.17 CB7.3 Bottom
6.6.17 CB3.3C Bottom
6.6.17 CB4.4 Bottom
6.6.17 CB5.3-1 Bottom
6.15.17 CB6.2 Bottom
6.5.17 CB7.3 Bottom
7.10.17 CB3.3C Bottom
7.10.17 CB4.2C Bottom
7.10.17 CB4.3C Bottom
7.10.17 CB4.4 Bottom
7.10.17 CB7.3 Bottom
8.31.17 CB3.3C Bottom
8.28.17 CB4.4 Bottom
8.28.17 CB5.3-1 Bottom
8.28.17 CB6.2 Bottom
4.12.17 CB3.3C Surface
4.10.17 CB5.3 Surface
4.10.17 CB6.2 Surface
4.10.17 CB7.3 Surface
6.6.17 CB3.3C Surface
6.6.17 CB4.4 Surface
6.6.17 CB5.3 Surface
6.5.17 CB7.3 Surface
7.10.17 CB3.3C Surface
7.10.17 CB4.4 Surface
7.10.17 CB7.3 Surface
8.31.17 CB3.3C Surface
8.28.17 CB4.4 Surface
8.30.17 CB5.3 Surface
8.28.17 CB6.2 Surface
8.28.17 CB7.3 Surface
B
Relative Abundance of 16S rRNA gene reads
100%
90%
80%
70%
60%
50%
40%
30%
20%
10%
0%
4.12.17 CB3.3C Bottom
4.10.2017 CB4.4 Bottom
4.10.17 CB6.2 Bottom
4.10.17 CB7.3 Bottom
6.6.17 CB3.3C Bottom
6.6.17 CB4.4 Bottom
6.6.17 CB5.3-1 Bottom
6.15.17 CB6.2 Bottom
6.5.17 CB7.3 Bottom
7.10.17 CB3.3C Bottom
7.10.17 CB4.2C Bottom
7.10.17 CB4.3C Bottom
7.10.17 CB4.4 Bottom
7.10.17 CB7.3 Bottom
8.31.17 CB3.3C Bottom
8.28.17 CB4.4 Bottom
8.28.17 CB5.3-1 Bottom
8.28.17 CB6.2 Bottom
4.12.17 CB3.3C Surface
4.10.17 CB5.3 Surface
4.10.17 CB6.2 Surface
4.10.17 CB7.3 Surface
6.6.17 CB3.3C Surface
6.6.17 CB4.4 Surface
6.6.17 CB5.3 Surface
6.5.17 CB7.3 Surface
7.10.17 CB3.3C Surface
7.10.17 CB4.4 Surface
7.10.17 CB7.3 Surface
8.31.17 CB3.3C Surface
8.28.17 CB4.4 Surface
8.30.17 CB5.3 Surface
8.28.17 CB6.2 Surface
8.28.17 CB7.3 Surface
Alphaproteobacteria
Actinobacteria
Gammaproteobacteria
Betaproteobacteria
Bacteroidetes
Cyanobacteria
Planctomycetes
Deltaproteobacteria
Verrucomicrobia
Clostridia
Bacilli
Chloroflexi
Other Proteobacteria
Thaumarchaeota
Others

**Figure S4**. Comparison between the relative abundance of the 14 most abundant phyla (and sub-phyla for Proteobacteria) between metagenomics and 16S rRNA gene datasets across 34 overlapping environmental samples. Sample names on x-axis are sorted by water column position (Top or Bottom), month (April, June, July and August), and station. Color legend at the bottom of the figure represent the same taxonomic groups in both A and B. (A) Relative abundance of taxa in metagenomic dataset in this analysis. Taxonomic groups are calculated as a percent of classified reads, and unclassified reads are excluded from this analysis. "Others" include all other classified taxa not explicitly listed. Photosynthetic algae are grouped with Cyanobacteria, since chloroplasts 16S rRNA gene amplify with the 16S rRNA gene primers used, so are included in the comparison. Other metagenomic reads classified as Animal or Virus are excluded from the analysis. (B) Relative abundance of taxa in 16S rRNA gene dataset from Arora-Williams et al. 2022. Relative abundance is calculated from all reads, and other includes both classified and unclassified 16S rRNA gene sequences. Chloroplast 16S rRNA gene sequences are included in the Cyanobacteria classification.

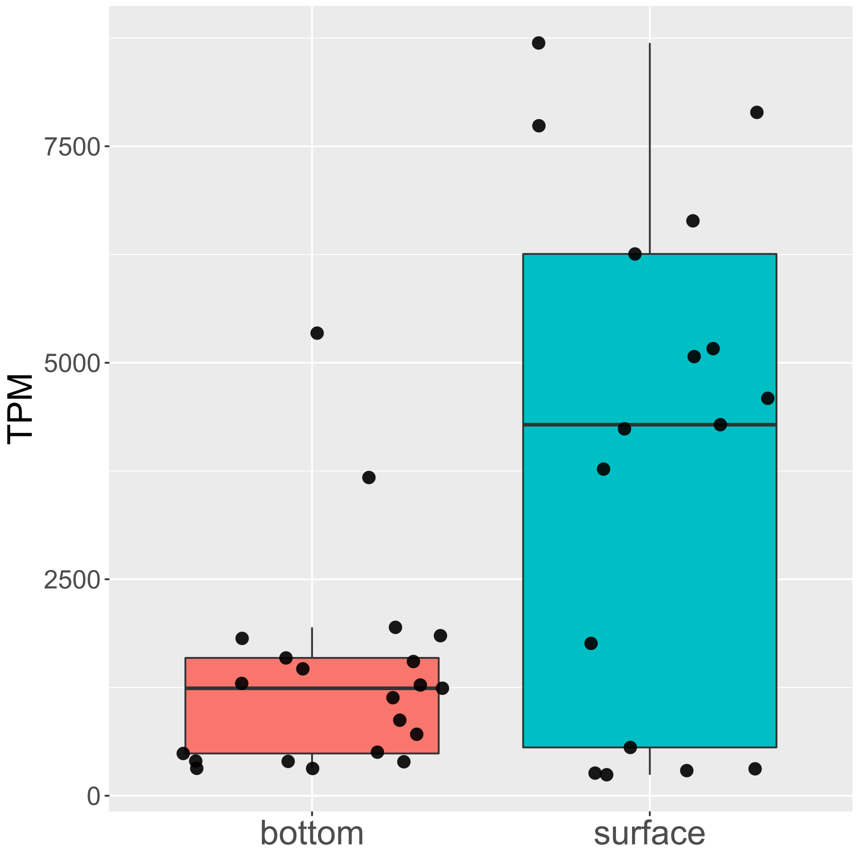


**Figure S5.** Differences in the relative abundance of Cyanobacteria between surface and bottom water samples. Relative abundance of reads classified as Cyanobacteria are plotted as box and whisker plots with all points displayed.

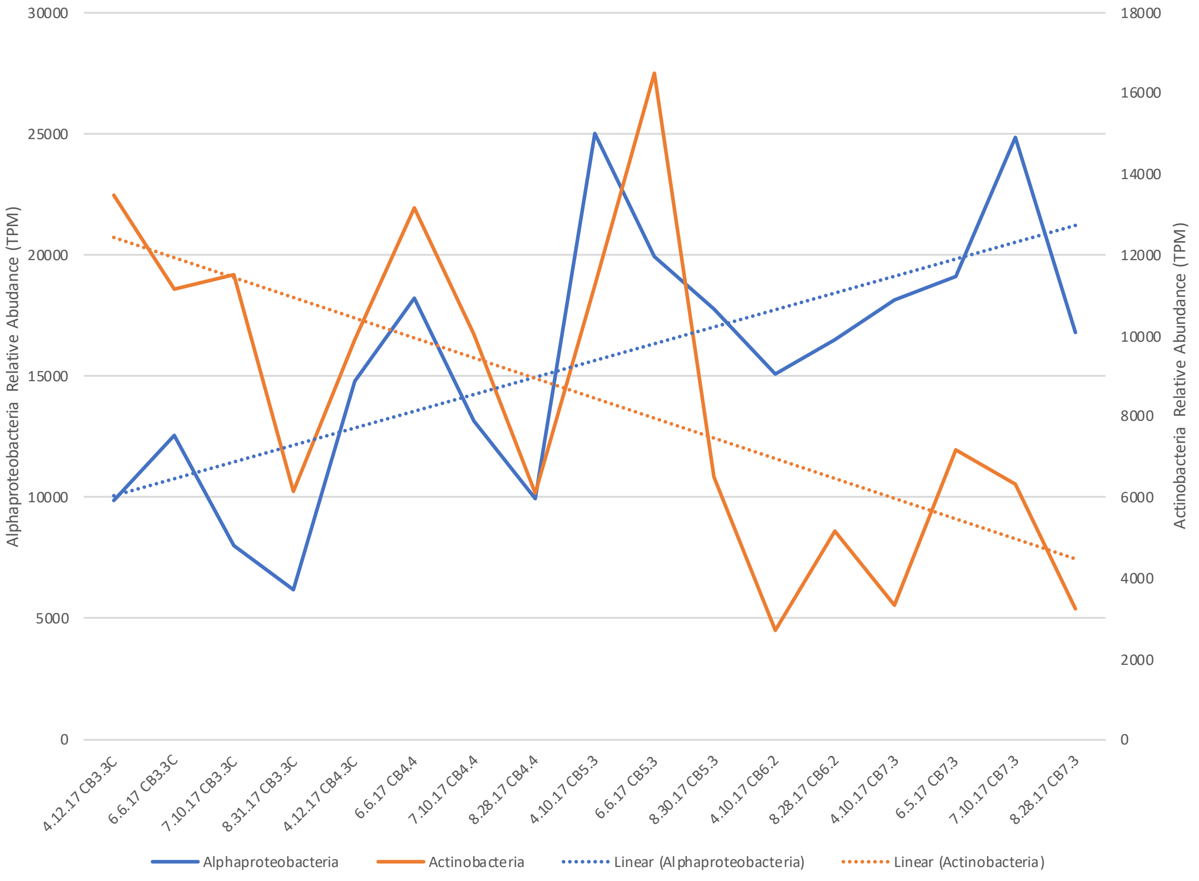


**Figure S6**. Distribution of Actinobacteria and Alphaproteobacteria across stations and time in the surface samples. Samples are arrayed from fresher (left, e.g. station CB3.3C) to marine (right, e.g. station CB7.3) and over time (4/12/17 to 8/31/17). Linear trend lines are shown for illustrative purposes only, to highlight general trend of decreasing Actinobacteria and increasing Alphaproteobacteria relative abundance towards the mouth of the Bay (left to right).

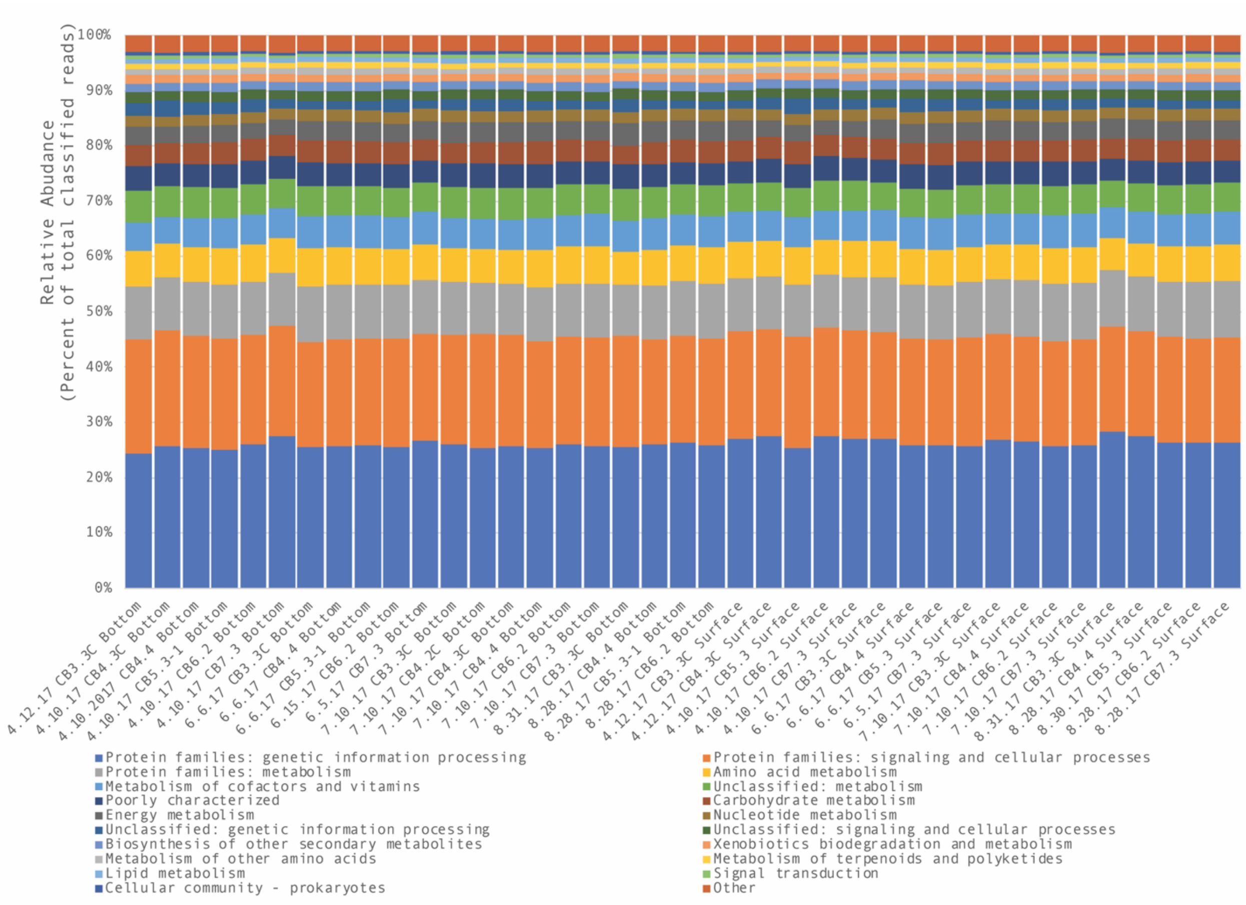


**Figure S7.** Relative abundance of genes classified in KEGG across samples. Sampling stations (x-axis) are arrayed according to water column position (Bottom or Surface), month (April, June, July and August), and station and labeled accordingly. The relative abundance of KEGG categories within the sample (aggregated at the second highest level in the KEGG hierarchy) as a percent of the total classified reads (genes without KEGG IDs are excluded for clarity). Very little change is observed across samples when comparing all KEGG categories.

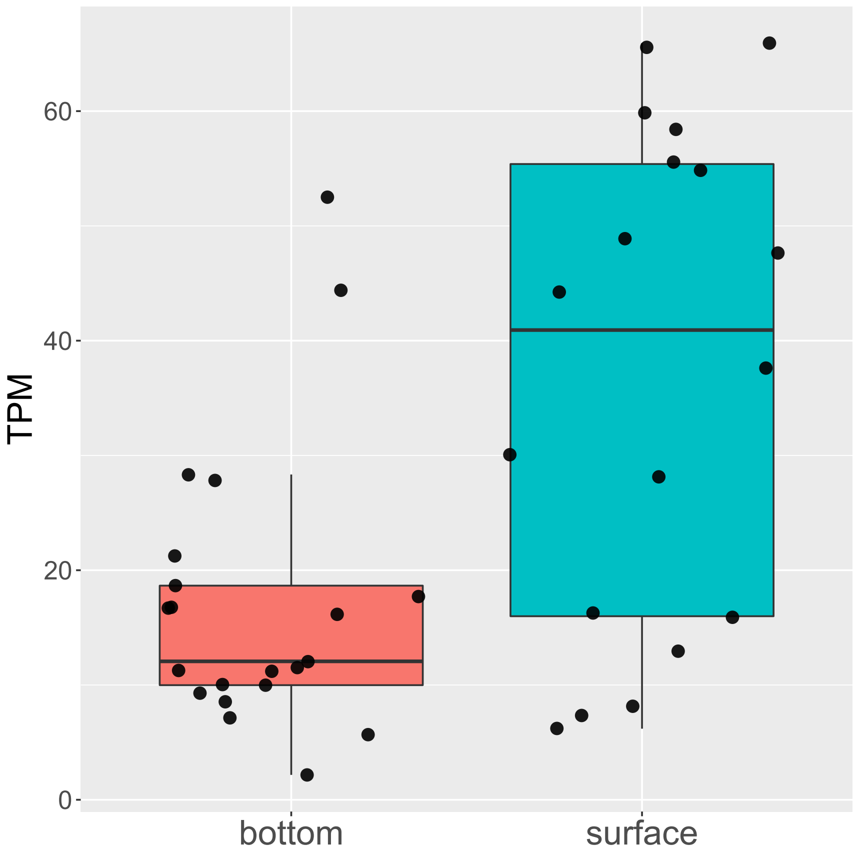


**Figure S8.** Difference between surface and bottom water samples in relative abundance of the photosystem II P680 reaction center D1 protein (*psbA*). All points are shown and the difference between surface and bottom water samples is statistically significant based on Student's T-Test ($p < 0.001$). Note that the bottom water sites include some locations in both April and August where mixing is relatively strong and where photosynthetic organisms would be expected to be mixed through the water column.

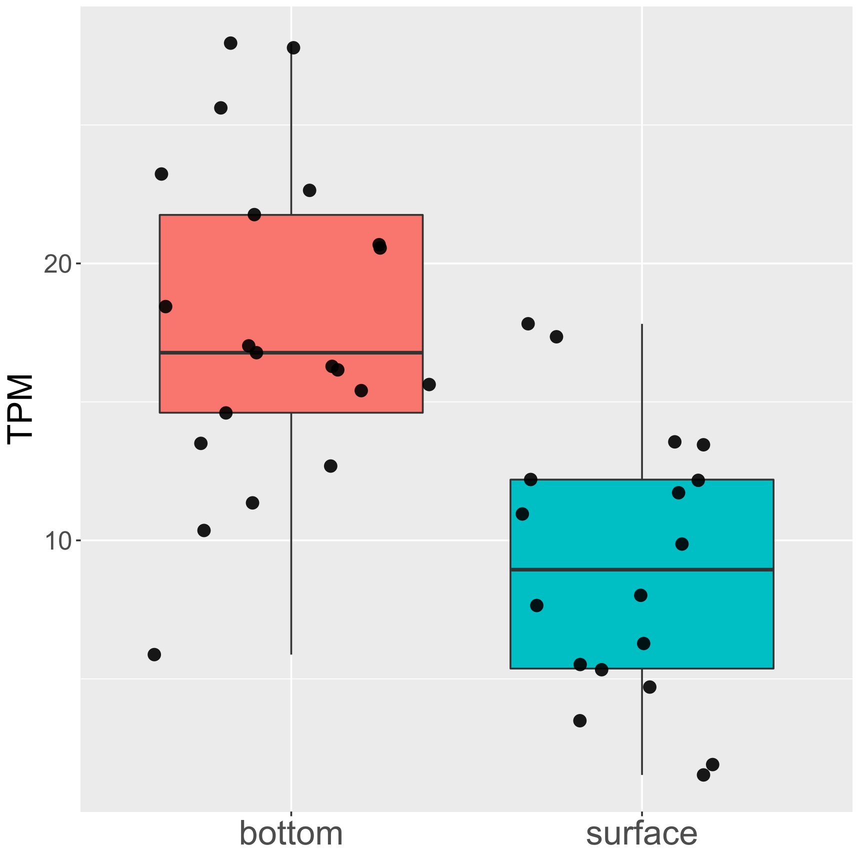


**Figure S9**. Difference between surface and bottom water samples in relative abundance of the adenylylsulfate reductase, subunit A (*aprA*). All points are shown and the difference between surface and bottom water samples is statistically significant based on Student's T-Test ($p < 0.001$).

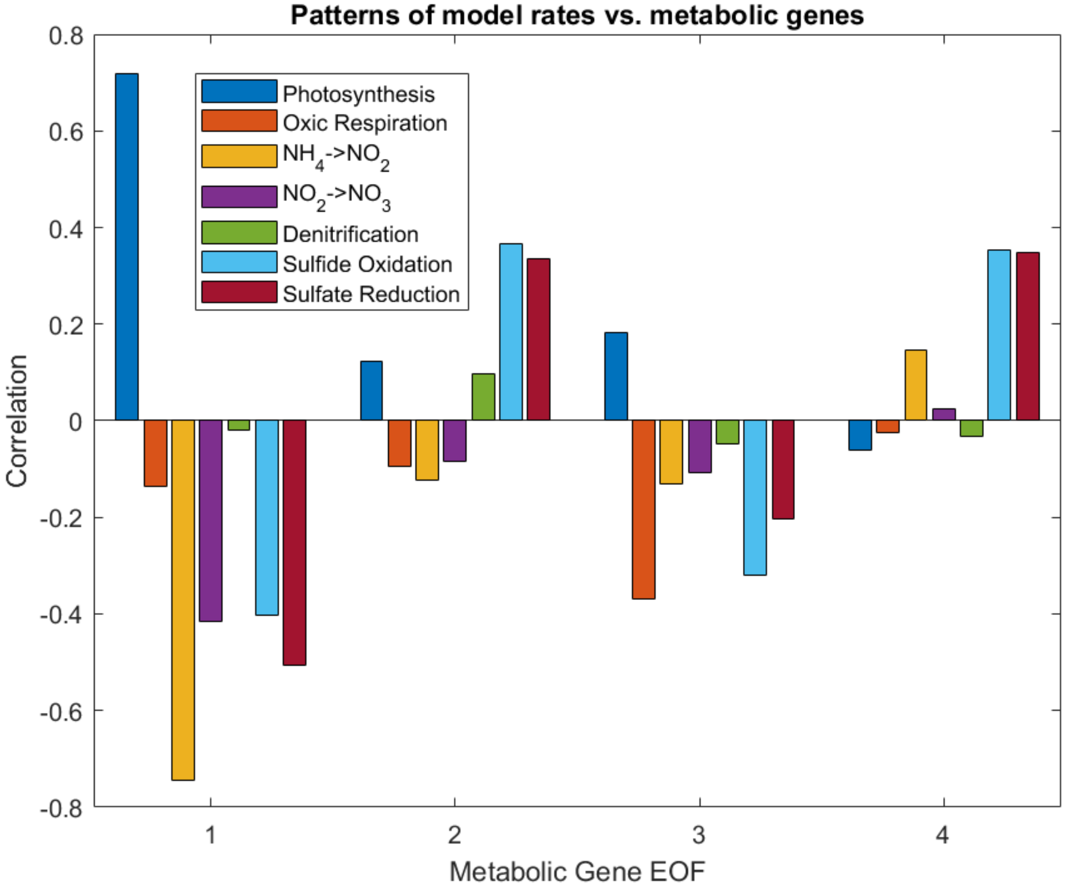


**Figure S10**. Correlation between the top four EOFs and the model predictions of key processes involved in carbon, nutrient and sulfur cycling. The EOFs also show relationships with the patterns of some, but not all biogeochemical rates predicted by the model.

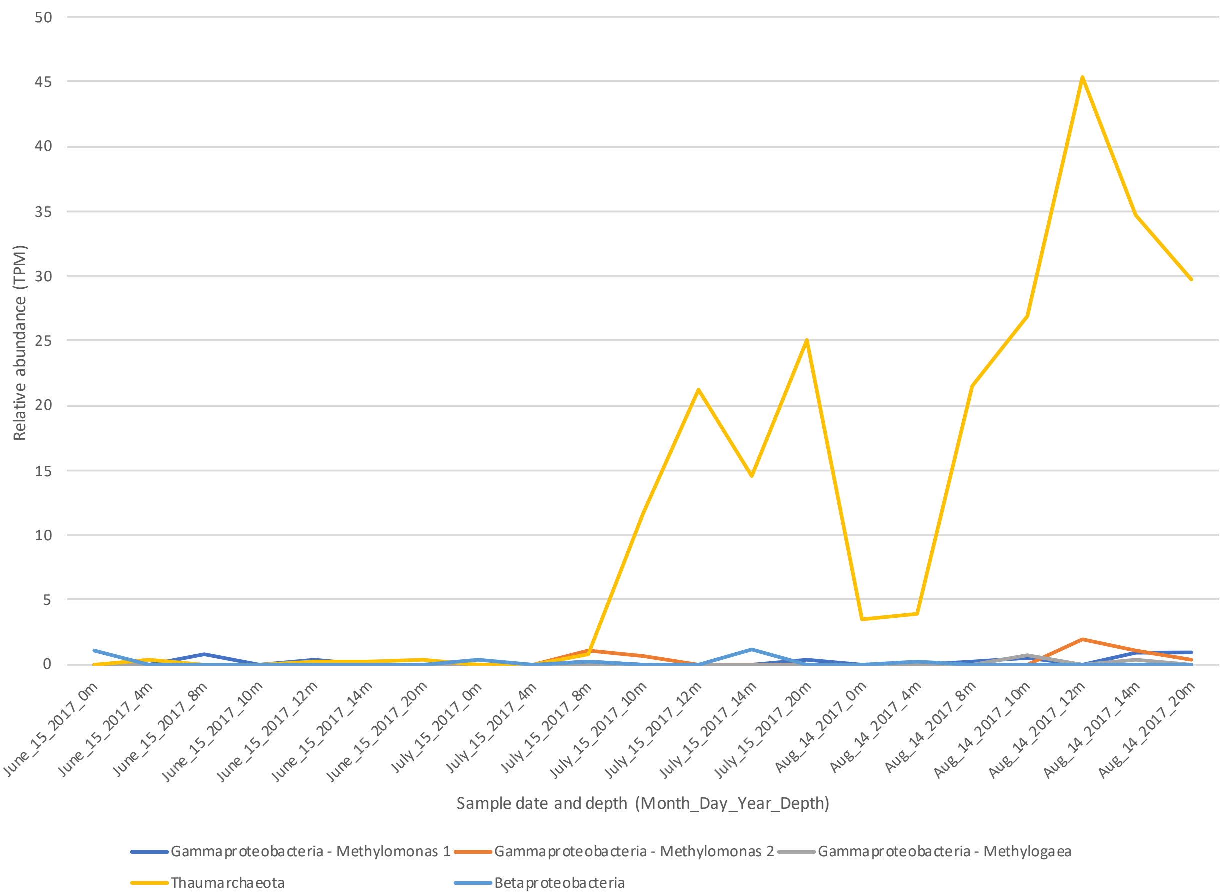


**Figure S11.** Relative abundance of *amoA* genes as a function of time and depth at one site (CB3.3C) in 2017. Sample dates and depths are displayed in the names on the x-axis, and are arrayed by date (June 15, July 15, and Aug 14), then depth (0, 4, 8, 10, 12, 14 and 20 m). The 20 meter depth is close to that used by the Maryland DNR for Mainstem sampling, demonstrating that abundance of *amoA* genes in the bottom sample is similar to those just below the thermocline. Archaeal *amoA* genes dominate at all positions in the water column in the late summer.

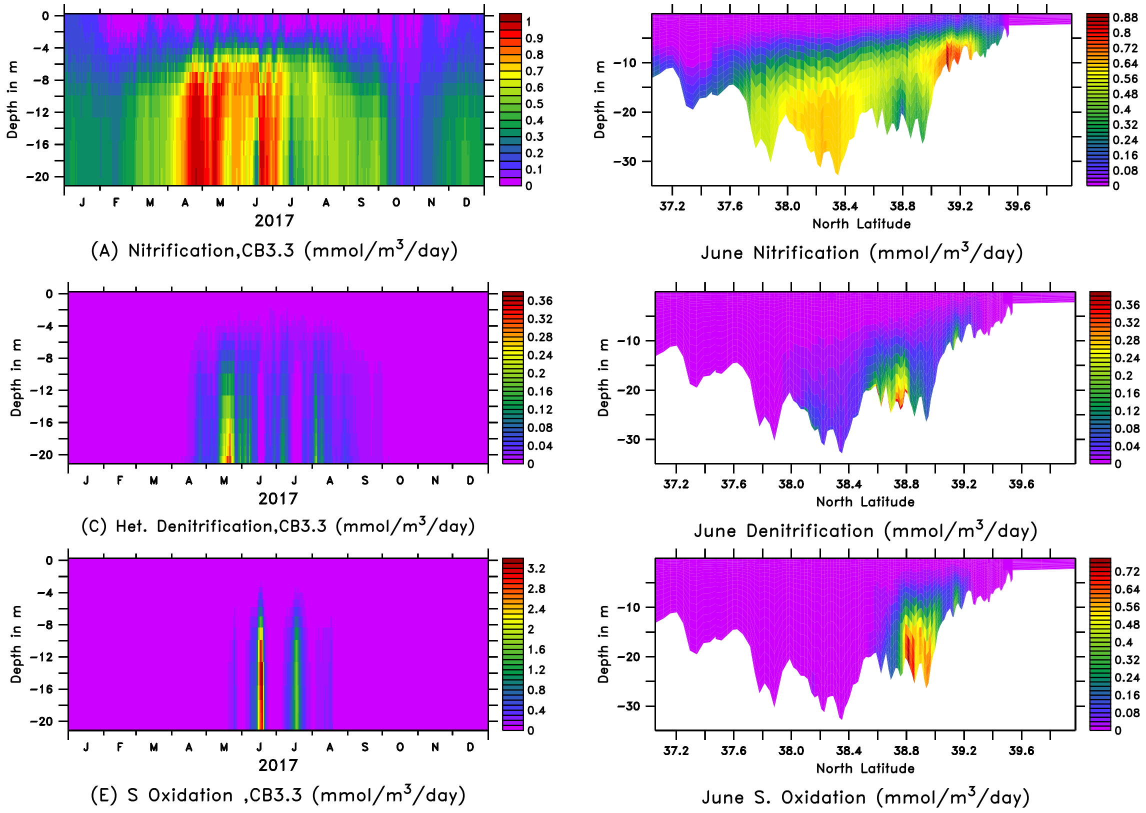


**Figure S12**. Predicted rates of nitrification, denitrification, and sulfide oxidation as a function of (a, c, e) depth and time at one site, or (b, d, f) depth and latitude in June. Colors indicate process rates (mmol $m^{-3}$ $day^{-1}$). Note that rates are similar between mid- column and bottom column, suggesting that sampling at the bottom of the water column should capture areas in the water column where these processes are expected to occur at some of the highest rates.

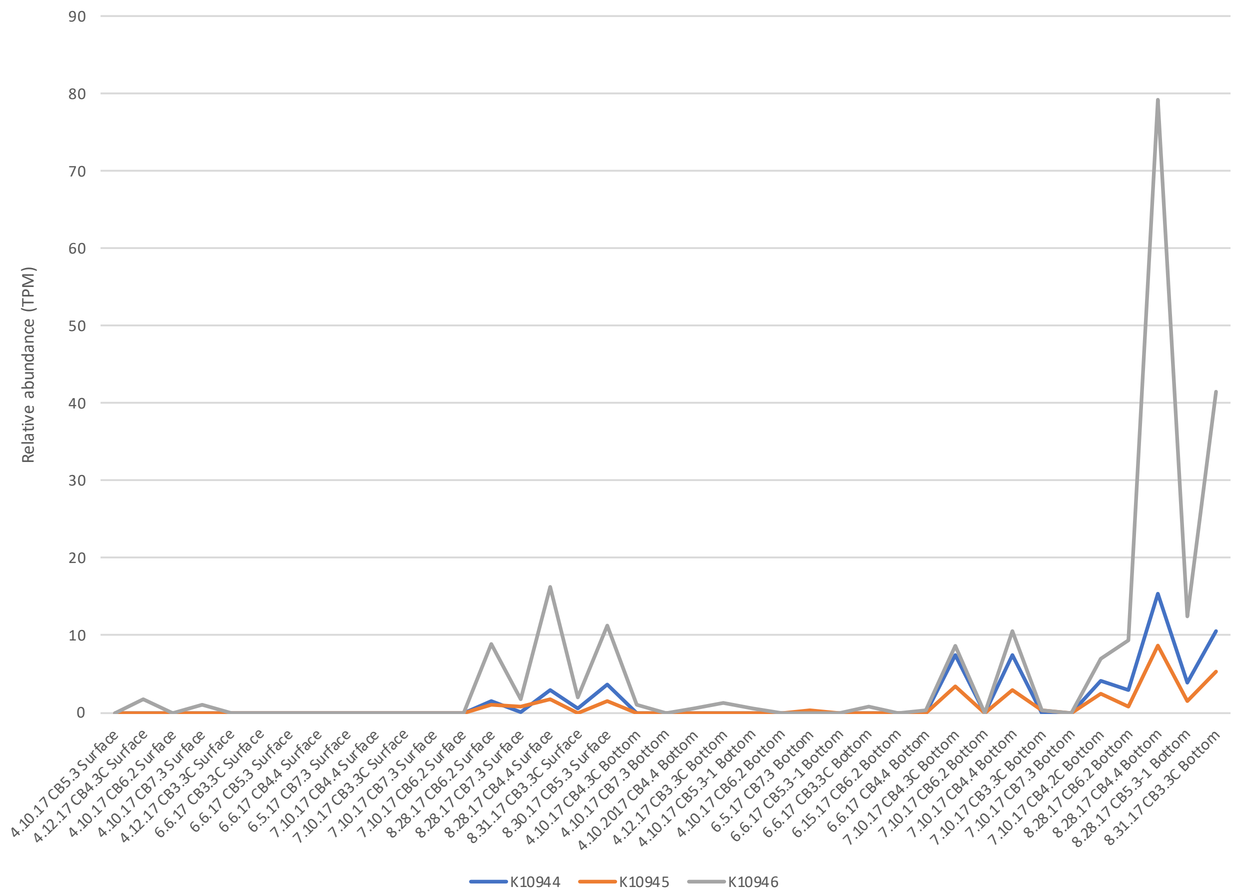


**Figure S13**. The relative abundance of *amoA* (blue, K10944), *amoB* (orange, K10945), and *amoC* (gray, K10966) across samples. Samples are listed on the x-axis by date (M.DD.YY), station (CB3.3C to CB7.3), and water column position (Surface or Bottom). *amoC* genes are much more abundant than *amoA* or *amoB* genes at the late summer sampling timepoint (8.28.17).

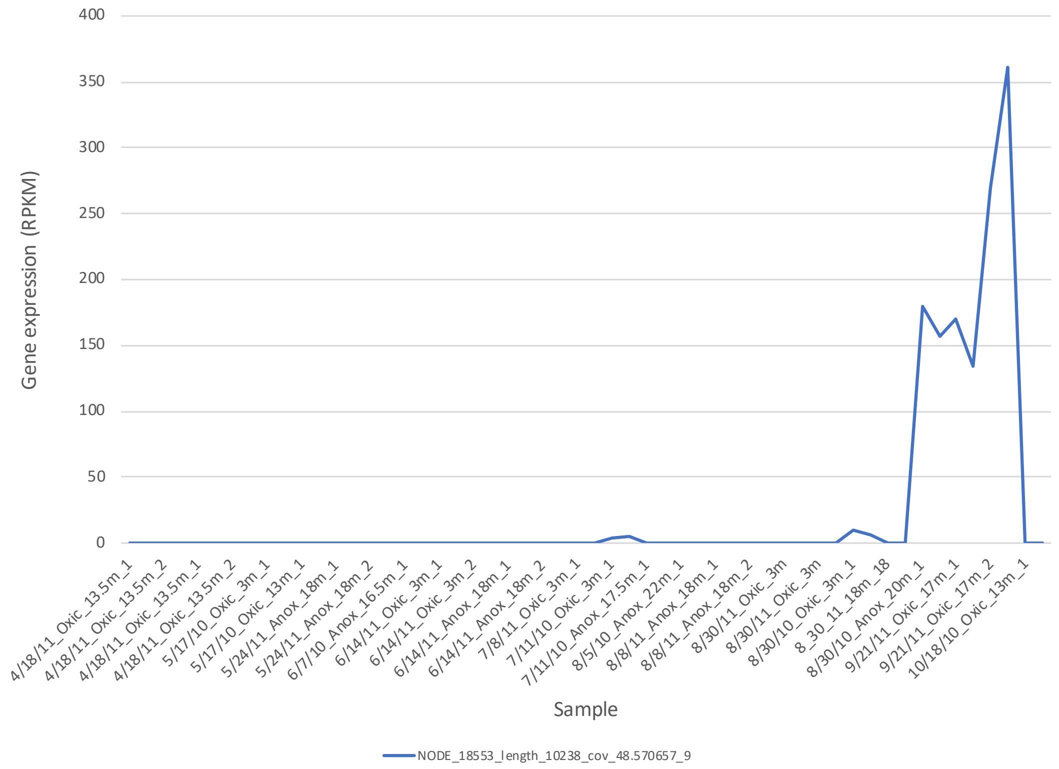


**Figure S14**. Expression of amoC gene that matches with 100% identity with a previously identified viral *amoC* gene in the Chesapeake Bay. Reads from metatranscriptomic dataset created by Hewson et al. from samples collected in 2010 and 2011, six to seven years prior to our sample collection. Samples were collected at one site near CB4.3 at different times of the year and depths above or below the oxycline, named according to date (mm/dd/yy), oxygen status (Anoxic or Oxic), depth (in meters), and replicate (1 or 2). This figure demonstrates the viral copy is expressed at the highest levels towards the end of the summer and into the fall in 2010 and 2011, suggestive of an active infection.

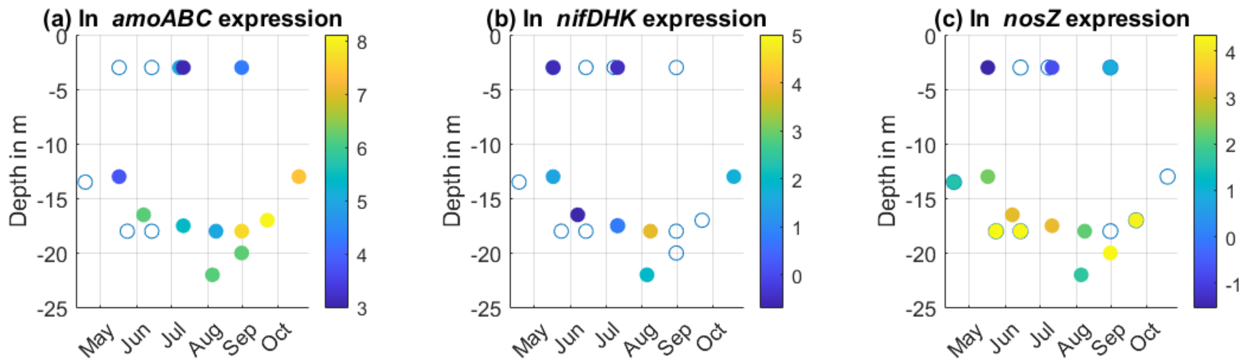


**Figure S15**. Expression of genes from 2010 and 2011, as determined by mapping reads from a previously published metatranscriptomic dataset (Hewson et al. 2014) to our metagenomic assembly. Points in space represent places and times when samples were collected. Dots are colored according to the log of the relative abundance of expression of these genes, in ln(reads per kilobase million, RPKM). Zero values represented with an open circle (pseudocounts were not added). a.) Expression of *amoABC* genes, ammonia monooxygenase subunit A, B and C. b.) Expression of *nifDHK* genes involved in nitrogen fixation. C.) Expression of nosZ genes involved in denitrification.

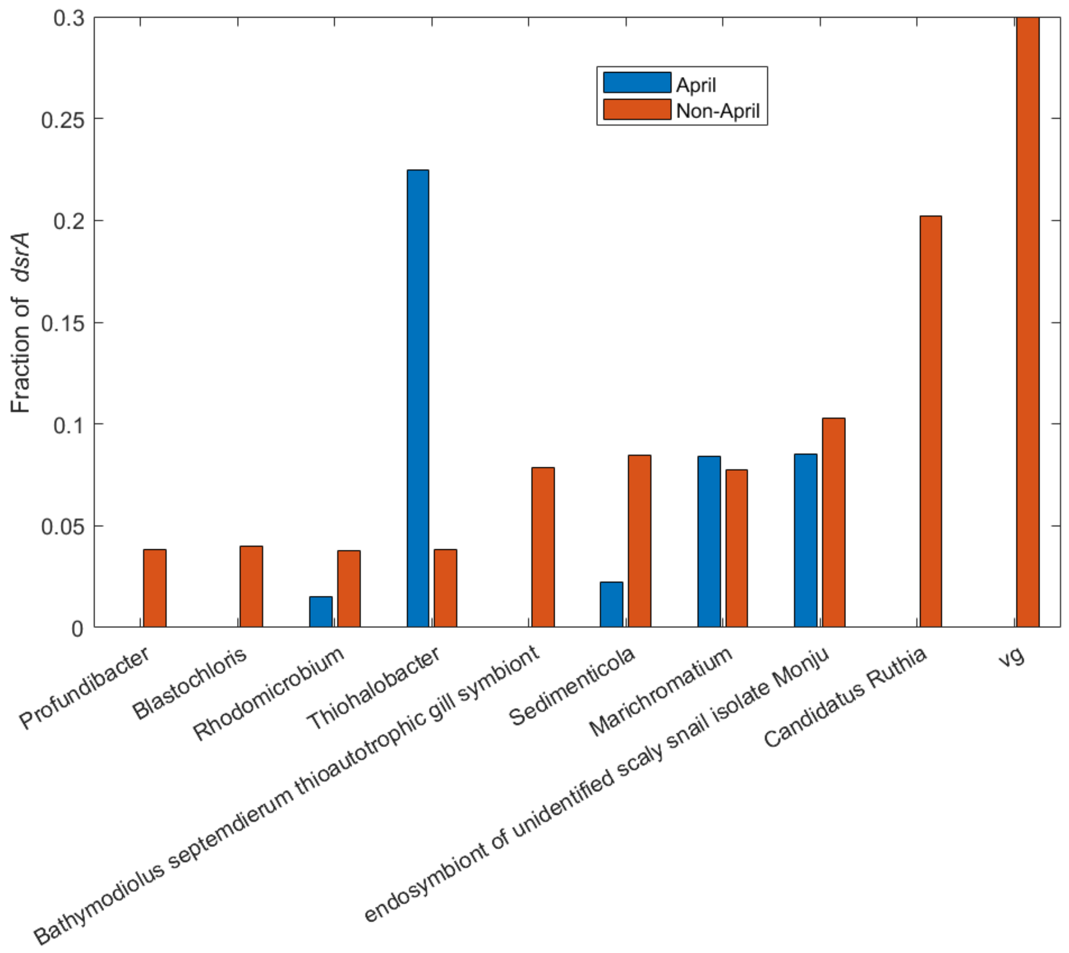


**Figure S16**: Differences in taxonomic classification of *dsrA* genes between April and the rest of the summer (non-April). The fraction of *dsrA* genes, weighted by relative abundance, classified according to the top 10 most abundant taxonomic groups (x-axis). Names associated with KEGG genomes are listed, including vg (Viruses) and two symbionts of invertebrates.
